# Regulatory start-stop elements in 5’ untranslated regions pervasively modulate translation

**DOI:** 10.1101/2021.07.26.453809

**Authors:** Justin Rendleman, Solomon Haizel, Shaohuan Wu, Junjie Liu, Xinyi Ge, Huijing Zou, Mahabub Pasha Mohammad, Matthew Pressler, Shuvadeep Maity, Vladislava Hronová, Zhaofeng Gao, Anna Herrmannová, Amy Lei, Kristina Allgoewer, Daniel Sultanov, Will Edward Hinckley, Ziyue Cheng, Lauren Shelby, Krzysztof J. Szkop, Ivan Topisirovic, Ola Larsson, Maria Hatzoglou, Leoš Shivaya Valášek, Christine Vogel

**Author notes:** Co-last authors (Leoš Shivaya Valášek, Christine Vogel). Co-first authors (Justin Rendleman, Solomon Haizel).

## Abstract

Sequence elements within the 5’ untranslated region (UTR) of eukaryotic genes, e.g. upstream open reading frames (uORFs), control translation of eukaryotic genes. We describe an element consisting of a start codon immediately followed by a stop codon which is distinct from uORFs in the lack of an elongation step. Start-stops have been described for specific cases, but their widespread impact has been overlooked. Start-stop elements occur in the 5’UTR of 1, 417 human genes and are more often occupied with a ribosome than canonical uORFs or control sequences. Start-stops efficiently halt ribosomes without evidence for accelerated RNA turnover, therefore acting as a barrier for the scanning of the small ribosomal subunit and repressing downstream translation. Our results suggest a model by which the ribosome undergoes repeated cycles of termination and partial ribosomal recycling, during which the large subunit detaches, but the 40S subunit with the Met-tRNA_i_^Met^ remains associated with the mRNA to be rejoined by the 60S subunit. Start-stop elements occur in many transcription factors and signaling genes, and affect cellular fate via different routes. We investigate the start-stop element in several genes, i.e. *MORF4L1*, *SLC39A1*, and *PSPC1*, and in more detail in *ATF4*.

## Introduction

The majority of mRNA translation in eukaryotes occurs in a 5’ cap-dependent manner. First, the 43S preinitiation complex (PIC) is recruited to the 5’ mRNA cap with the help of the eukaryotic translation initiation factors 4F (eIF4F) and eIF3. Subsequently, these factors together with other eIFs facilitate the scanning of the 48S PIC along the 5’ untranslated region (UTR) of the mRNA until the start codon is recognized and translation is initiated (1–3). Hence, the scanning process naturally compels the 48S PIC to read the genetic information encoded in the 5’ UTR, which includes a number of translation control elements that affect protein production, often in a condition-dependent manner (4). Despite decades of extensive research, understanding of the regulatory code embedded in the 5’ UTR is still incomplete, largely due to the sheer number of elements and plausible regulatory outcomes.

Upstream open reading frames (uORFs) are one of the regulatory elements frequently found in 5’ UTRs, which occur in approximately half of human mRNAs (5). Functional uORFs undergo standard translation, including initiation, elongation producing the uORF peptide, and termination. uORFs often act to repress translation of downstream coding regions (5), although exceptions exist, where they act as translation reinitiation-inducible elements (6–12). There is also a special class of uORF-like elements composed of an AUG immediately followed by a stop codon. Because it inherently excludes an elongation step, it does not represent a *bona fide* open reading frame, and we refer to this element as start-stop.

Recent literature has revealed examples of regulatory start-stops, such as in the 5’ UTR of *Arabidopsis thaliana* genes, including NIP5;1 (13). NIP5;1 encodes a boric acid channel, and the start-stop appears to stall 80S ribosomes in a boron-dependent manner, triggering mRNA degradation and thereby preventing expression of the gene. A similar, boron-dependent effect on translation was observed for start-stop sequences in other plant and yeast genes (14–16). Most recent analyses suggested a highly specialized mechanism in which the start-stop halts the ribosome during boron stress, whereby boron interferes with correct termination of the ribosome (17).

In mammals, a start-stop has been described in the mRNA leader of *HOXA11* encoding an embryogenesis-promoting transcription factor (18). In this gene, the start-stop was thought to interact with a highly stable stem-loop to inhibit *HOXA11* mRNA translation. However, other work suggests that the stem-loop is sufficient to stall 80S ribosomes and the start-stop might not be transcribed as part of the 5’UTR examined originally (19). A start-stop also resides upstream of two major regulatory uORFs within the 5’UTR of the gene *ATF4*, encoding a master transcription factor of the integrative stress response (12, 20–23) - it is conserved across mammals but absent from mouse. The *ATF4* start-stop’s function is thought to depend on the non-canonical initiation factors DENR and MCTS1, as DENR knockout partially reduced the overall stress-induced upregulation of *ATF4* mRNA translation (21). The same work also suggested that DENR/MCTS1 in general are critical for re-initiation after start-stop elements, i.e. for the translation of the downstream open reading frames. More recent work in yeast does not support this view (24).

Therefore, the general role of start-stops in mammalian genes is not clear, even though they occur in hundreds of human genes (21) and are significantly enriched in the 5’UTRs of genes that are most intolerant to loss-of-function mutations (25). As start-stops are distinct from uORFs in terms of the omission of the elongation step, we hypothesized that they must affect ribosome processivity through a non-canonical mechanism that involves termination without a single elongation cycle. Here, we show that start-stop elements indeed stall ribosomes and suppress downstream translation more efficiently than even the smallest canonical uORF. They occur in >1, 400 human genes, including many signaling molecules and transcription factors, are highly conserved, and do not trigger mRNA degradation (unlike uORFs). Start-stops represent an important class of regulatory elements that possibly act through repeat cycles of termination and partial ribosome recycling.

## Methods

### Ribosome occupancy across elements and datasets

We downloaded GENCODE v43 human gene annotation file from the UCSC Table Browser (https://genome.ucsc.edu/cgi-bin/hgTables). When a gene had multiple principal transcripts, we selected the transcript with the longest 5’ UTR. To extract the 5’ UTRs, which include start-stop sequences (‘AUGUGA’, ‘AUGUAA’, or ‘AUGUAG’), as well as control sequences (‘AUUGGA’, ‘AUUGAA’, or ‘AUUGAG’), we used a customized R script. In total, we obtained 1, 417 and 1, 837 genes containing start-stop and control sequences in their 5’ UTRs, respectively (**Table S1**).

We downloaded ribosome profiling and RNA-seq data collected in HeLa cells from the Gene Expression Omnibus (GSE113171), i.e. SRA files (study ID SRP140470). In this experiment, HeLa cells were treated with 0.5ug/mL tunicamycin for 0, 1, 4, or 8 hours, and prior to harvest cell growth media was removed and cells were washed with fresh media supplemented with 0.1 mg/mL cycloheximide for five minutes (26). We converted the SRA files to fastq format and performed trimming using the cutadapt algorithm (version 1.12) to remove low quality bases at the 3’ and 5’ ends of reads (-q 30, 30), and to discard reads shorter than 25 nt (-m 25)(27). Trimmed fastq files were then aligned to the human genome (NCBI GRCh38) using the HISAT2 aligner (version 2.0.5), with a seed value set to 23 (--seed 23), and no soft clipping of reads (--no-softclip)(28).

We examined the quality of the ribosome footprint data using the RiboTaper analysis pipeline (29, 30). The analysis was based on the observation that genuine ribosome footprint reads typically exhibit a distinct 3-nt periodicity within coding regions. We confirmed this pattern in reads with length ranging from 28-30 nt in the Hela dataset. Therefore, we counted ribosome footprints when the length of the read fell within the 28-30 nt range and its starting position was exactly 12 nt upstream of the AUG or AUU in start-stop or control sequences, respectively (P-site position). We excluded reads with other lengths or positions from our analysis. We used htseq-count to assign reads to genes and to start-stops (31). GTF files were modified to count reads mapping to start-stops in the 5’ UTR by excluding any overlap with the coding sequence (CDS) and requiring a 12 nt distance from the start codon. We required the start-stop’s P-site to be located completely within the 5’ UTR, but allowed for partial overlap of the 3’ end of reads with the CDS to capture ribosomes approaching the start site.

To generate the heatmap displaying fractional occupancies of all P-site/A-site combinations, we compiled ribosome footprint data for all ribosome reads in the 5’UTR (**Table S2**). The resulting data was then clustered using the Perseus software (version 1.5.0.15), using the Euclidean clustering method (32).

We repeated the analysis with ribosome profiling and RNA-seq data from HEK293 cells (30), mouse primary keratinocytes (33)(**Table S5**), and *Saccharomyces cerevisiae* (34)(**Table S6**). For the HEK293 data, we followed the same processing steps as described above. Mouse sequencing data was first trimmed for the adapter sequence AGATCGGAAGAGCACACGTCT and then aligned to the NCBI GRCm38 reference genome. Yeast sequencing reads were trimmed with cutadapt to remove polyA adapter sequences, and subsequently aligned to the NCBI *S. cerevisiae* build3.1 reference genome.

We expanded the analysis to a dataset from Hela cells generated without the use of translation inhibitors that could potentially interfere with ribosome positioning and footprint length (35) and compared ribosome occupancy on start-stop elements, minimal uORFs, and control sequences. We defined minimal uORFs as sequences which started with “AUG” and ended with “UGA”, “UAA”, or “UAG”, and had one of the 59 sense codons in the middle (excluding AUG as a middle codon). We removed the adaptor and random sequences from the raw reads with Cutadapt v3.1 (27). Subsequently, we aligned the trimmed reads against the human genome GRCh38.p10 with STAR v2.7.6a (27, 36), with similar parameter settings to those described in reference (35): --outFilterIntronMotifs RemoveNoncanonicalUnannotated --outFilterMultimapNmax 1 --outFilterMismatchNoverLmax 0.1 --outSAMtype BAM SortedByCoordinate --outSAMunmapped Within --outSAMattributes Standard. We constructed lists of start-stop, control, and uORF containing genes as described above, using GENCODE v43 human gene annotation file (https://genome.ucsc.edu/cgi-bin/hgTables). We then used a customized R script to count reads across all positions relative to the start and stop sites of the start-stop sequences and of the uORFs, and the first and last codons in the control sequence. This analysis was conducted with a 40 nt window surrounding each element.

### Characteristics of start-stop containing genes

We compiled a high-confidence core set for human data using the following criteria. Occupied start-stop and control sequences were required to have 3 or more RPF reads in both the HeLa and HEK293 datasets; unoccupied start-stop and control sequences were required to have 0 RPF reads, but at least 6 RNA reads in both HeLa and HEK293 datasets. In addition, we manually quality controlled the resulting sets (**Table S3**). These stringent criteria resulted in 88 and 7 occupied start-stop and control sequences, respectively. Additionally, the core set comprised 21 and 85 unoccupied start-stop and control sequences.

To estimate the number of ribosomes involved in translating the mRNA transcript of each corresponding gene, we calculated the translation efficiency as log_2_(RPF/RNA) within the CDS region of the gene. Statistical analysis was performed using a two-sample t-test.

To calculate evolutionary conservation, we downloaded conservation scores from the UCSC Genome Table Browser data retrieval tool (http://genome.ucsc.edu). For each element (start-stop-occupied, start-stop-unoccupied, control-occupied, control-unoccupied), phyloP evolutionary conservation scores from 100 vertebrate species were averaged across the hexamer sequence and statistical comparison of the start-stop-occupied elements with the other three groups was performed by two-sample t-tests, after exclusion of outliers (**Table S4**).

To calculate nucleotide frequencies surrounding specific elements, we extracted sequences surrounding start-stops from transcript annotations. Relative nucleotide frequencies were then calculated for ±15 nt from the beginning and end of each start-stop for comparison.

For analysis of enriched functions, we compiled lists of genes that contained occupied start-stop sequences in their 5’ UTRs for HeLa, HEK293, and mouse primary keratinocytes, using less stringent cutoffs: for occupancy, we required at least three RPFs and at least two RPFs on the start-stop in the human and mouse data, respectively (**Table S5**).

We analyzed the set of start-stop containing genes identified i HeLa, HEK293, and mouse primary keratinocytes for enriched functions using IPA (QIAGEN Inc., https://www.qiagenbioinformatics.com/products/ingenuity-pathway-analysis)(37)(**Table S8**). We analyzed the total list of 1, 417 start-stop containing genes for enriched functions using the NIH DAVID annotation tool (https://david.ncifcrf.gov/)(**Table S9**). P-values were adjusted for multiple hypothesis testing using the Benjamini-Hochberg correction (38).

### Assessing the effect of start-stop elements on translation and mRNA stability

We used the dual-luciferase psiCHECK2 system (Promega) following the manufacturer’s protocol. The psiCHECK2 vector expresses both firefly luciferase and renilla luciferase, allowing for an internal control within the same plasmid. HEK293T or HeLa cells were seeded in a 24-well plate at a density of 4 x 10^4^ cells in 1 mL of Dulbecco’s Modified Eagle Medium (DMEM) growth media supplemented with 10% fetal bovine serum (FBS, Thermo Fisher) and penicillin/streptomycin (Gibco). The cells were then incubated at 37 °C and 5% CO_2_ until they reached approximately 70% confluency. We transfected cells using 1 ug of reporter plasmid and TurboFect™ (Thermo Fisher) transfection reagent, following the manufacturer’s protocol. Subsequently, we used the Dual-Glo® luciferase assay kit (Promega) to measure the activities of both firefly and renilla luciferase in the cell lysate harvested from each well of the 24-well plate, as per the manufacturer’s instructions.

To test the function of the start-stops in MORF4L1 (NCBI Reference Sequence: NM_006791.4), SLC39A1 (NCBI Reference Sequence: NM_001271958.2), and PSPC1 (NCBI Reference Sequence: NM_001354909.2), we inserted the 5’ UTRs of these respective human genes upstream of the renilla reporter gene (hRluc) at the NheI site of the psiCHECK2 plasmid. Start-stop positions and mutants were as follows: MORF4L1 - start-stop 108 nucleotides upstream of the CDS; St-st Mut: AUGUAG->UUGUAG. SLC39A1 - start-stop 47 nucleotides upstream of the CDS; St-st Mut: AUGUAG->UUGUAG. PSPC1 - start-stop 75 nucleotides upstream of the CDS; St-st Mut: AUGUAG->AUUGAG.

To examine the role of the start-stop in a gain-of-function construct, we used the 5’ UTR of β-Actin (ACTB, NCBI Reference Sequence: NM_001101.5). It has a length of 84 nucleotides. The start-stop or uORF were positioned either 77 or 30 nucleotides upstream of the luciferase CDS to mark ‘near cap’ or ‘near CDS’ positions, respectively.

Sequences were mutated using site-directed mutagenesis with primers as shown in **Table S11.** All transfections were performed in triplicate as biological replicate, and both firefly and renilla luciferase activities were measured in triplicate as well. We then calculated relative luciferase activity by dividing the averaged reporter luciferase activity by the control luciferase activity. This value was further normalized by relating all measurements to that of the respective wild-type 5’ UTR. Statistical analysis was conducted using two-tailed, paired t-tests, where samples were paired by the day of the biological replicate.

To estimate mRNA stability, HEK293T or Hela cells were transfected with 1ug of the respective plasmid. Total RNA was isolated from the cells using the RNeasy kits (Qiagen) according to the manufacturer’s instructions. Reverse Transcription of cDNA was performed using SuperScript III Reverse Transcriptase (Thermo Fisher Scientific).

Quantitative RT-PCR was carried out to measure firefly (hluc) and renilla (hRluc) RNA levels using KAPA SYBR FAST qPCR master mix (Kapa Biosystems) with the following program: preincubation at 95 °C for 3 minutes followed by 40 cycles of 95 °C for 10 seconds, 60 °C for 30 seconds, and 72 °C for 30 second. We then analyzed the samples with the Roche LightCycler 480 system. **Table S11** lists the primer sequences for the RT-PCR.

### Comparing start-stops to minimal uORFs

We predicted 9 nucleotide uORF (“minimal uORFs”) sequences as we predicted start-stop and control sequences (see above), searching for AUG and in-frame stop codons, with a single codon in the middle. To compare start-stops and minimal uORF, we examined a published large-scale dataset (39). In their analysis, the authors constructed a library of randomized 50-nucleotide 5’ UTR oligomers which were fused to the eGFP CDS. They then analyzed the effect on translation by separating sequences by polysome profiling and sequencing the respective fractions (mono-, di, polysome fraction). We compared the placement of sequences containing start-stops, minimal uORFs and control sequences as defined above. We next extracted sequences for the same three elements from a large-scale screen on determinants of transcript stability (40).

To examine the ribosome footprint lengths as a measure of A-site occupancy, we extracted reads positioned to the AUG or stop codon of the start stop, the uORF, or CDS, or the first and last codon of control sequences from the datasets by Wu et al. which did not employ cycloheximide or other translation inhibitors (35). We then counted short (20-22 nt) and long (27-30 nt) footprint reads separately. We repeated this analysis for ribosomes at the stop codon of the uORF (A-site) and for the AUG and stop codons of the main CDS. By calculating the average ratios of the short read counts to long read counts across different positions, we obtained an estimate of the occupancy status of the A-site-containing ribosomes.

To analyze the ratio of 40S/80S ribosome footprints, we downloaded two published datasets (12, 21), labeled as Dataset 1 and 2, respectively. These methods measure footprints of the 48S PIC (40S) subunits and also the fully assembled 80S subunit. We post-processed the data as described above and compared read mappings for start-stop elements, control sequences, and the start and stop codons of minimal uORFs and the CDS. We determined the dominant position and length ranges of reads at the elements of interest, i.e. the AUG of the start-stop, uORF, or CDS, the AUU for the control sequence, and the respective stop codons. For Dataset 1 (12), we counted 40S and 80S reads with a length between 30 and 36 nucleotides with the 5’ end of the read positioned to -14 to -12 nucleotides relative to the element of interest. For Dataset 2 (21), we counted 80S reads for lengths between 30 and 33 nucleotide positioned at -13 to -12 nucleotides, and 40S reads for lengths between 26 to 29 nucleotides positions at -10 to -9 nucleotides. We then aggregated counts across all genes in a metagene analysis and calculated the 40S/80S ratios.

### Analyzing the start-stop in ATF4

To examine ribosome density along the ATF4 5’ UTR and beginning of the CDS, we used samtools-depth algorithm (41) and published Hela data (26). Values were normalized based on library size, using size factors from the DESeq2 R package (42). To account for variation in ATF4 mRNA levels, we calculated the relative ribosome density for each sample as the ratio of normalized ribosome density divided by the normalized ATF4 mRNA abundance. These relative ribosome densities were then plotted along the ATF4 genomic coordinates (26).

We used Crispr-Cas9 based genome editing to modify the start-stop in ATF4 in the human HAP1 cell line. The HAP1 cell line is near-haploid which facilitates generation of a homozygous mutant at the start-stop site. To design Cas9 guide RNA targeting this site, we identified a PAM sequence (NGG) in the ATF4 5’ UTR region surrounding the start-stop site. To assess potential target sequences, we employed the Benchling CRISPR Guide RNA Design software (https://benchling.com). Among the options, we selected the guide target sequence CGCAGAGAAAACTACATCTG, which possessed an on-target score of 57.1 and an off-target score of 59.9 (43, 44).

We then used the lentiCRISPRv2 plasmid (AddGene 52961) as a one vector system for co-delivery of the hSpCas9 protein and the sgRNA; cloning of the target sequence into the plasmid was performed as previously described (44). Two oligonucleotides with overhangs were synthesized for cloning the selected sgRNA into the lentiCRISPRv2 plasmid (**Table S11**). The oligos were phosphorylated and annealed with T4 PNK (NEB) supplemented with 10mM ATP at 37 °C for 30 minutes, followed by 95 °C for 5 minutes, and a ramp from 95 °C to 25 °C at 5 °C/minute. Next, the plasmid backbone was digested and dephosphorylated using BsmBI restriction enzyme (NEB) and Quick CIP (NEB), with incubation at 55 °C for 1 hour and heat inactivation at 80 °C for 20 minutes. We performed gel purification using the QIAquick Gel Extraction kit (Qiagen), and ligated the linearized backbone to annealed oligos using Quick Ligase (NEB). Transformation into NEB Stable Competent *E. coli* bacteria (NEB) was performed according to manufacturer instructions. Single colonies were selected, and plasmid DNA was purified from overnight cultures using the QIAprep Spin Miniprep Kit (Qiagen). Proper annealing of sgRNA into the lentiCRISPRv2 backbone was confirmed by Sanger sequencing (Azenta) using the appropriate primer (**Table S11**).

HAP1 cells were cultured in Iscove’s Modified Dulbecco’s Medium (IMDM, Corning) supplemented with 10% fetal bovine serum (FBS, Thermo Fisher) at 37 °C and 5% CO_2_ until they reached approximately 40% confluency. The lentiCRISPRv2 plasmid containing the start-stop target sequence was transiently transfected into HAP1 cells using Lipofectamine 3000 (Thermo Fisher) and incubated for 24 hours. Cells were then passaged into a fresh growth medium supplemented with 0.4 ug/mL of puromycin for selection of positive transfection. After three days of growth in the selection medium, individual cells were isolated into separate wells of a 96-well plate using fluorescence-activated cell sorting (FACS). Clonal populations were subsequently grown out and PCR screening was conducted to confirm the presence of the desired band size within the ATF4 region before being sent for Sanger sequencing (Azenta) using primers as shown in **Table S10.** Three lines were selected for follow up based on the sequences at the start-stop in ATF4 5’ UTR: Start-stop^WT^ cells, which had no differences in sequence compared to the wild-type human genome, ΔStart-stop cells, in which the start-stop sequence was completely deleted, and Start-stop^TAGTAG^ cells, where the start codon was mutated to a stop codon (AUG->UAG). For stress induction experiments, cells were incubated with a final concentration of 1 uM thapsigargin.

We used Western blotting to measure ATF4 protein levels in the different HAP1 cell lines. To do so, we diluted ATF4 Rabbit monoclonal antibody (Cell Signaling) in a ratio of 1:1000 into a Clean Western Buffer with 5% BSA. Next, we washed the membranes three times in Clean Western Buffer for 5 minutes each and then incubated for 1 hour with Digital Anti-Rabbit secondary antibody (Kindle Biosciences) at 1:2000 in Clean Western Buffer with 5% BSA. Proteins were detected with the KwikQuant Imaging System (Kindle Biosciences). As a loading control detection, we used the same Western protocol, replacing the primary antibody with the β-Actin Rabbit antibody (Cell Signaling) after incubating the blots with stripping buffer (Thermo Fisher).Western blot images were processed with the ImageJ software to quantify protein levels for ATF4 and loading controls. We subtracted background values for each sample and divided ATF4 values by loading control values to account for variability in sample loading between wells. To allow for comparisons across the three replicate blots, these normalized ATF4 values were then transformed based on the percent of the maximum value from each blot (relative % of max).

To measure ATF4 transcript levels in the different HAP1 cell lines, we isolated total RNA from cells using the RNeasy kits (Qiagen) following the manufacturer’s protocol. cDNA was reverse-transcribed from RNA samples using SuperScript III Reverse Transcriptase (Thermo Scientific).

To estimate the decay of the ATF4 mRNA, we treated Start-stop^WT^ and ΔStart-stop cells with either RNA polymerase II inhibitor 5, 6-dichloro-1-β-ribofuranosyl-benzimidazole (+) or DMSO (-) and isolated RNA after 2 hours.

Quantitative RT-PCR was carried out using the KAPA SYBR FAST qPCR master mix (Kapa Biosystems) with samples pre-incubated at 95 °C for 3 minutes then 40 cycles of 95 °C for 10 seconds, 60 °C for 30 seconds, and 72 °C for 1 second, and analyzed with the Roche LightCycler 480 system. The RT-PCR primer sequences are listed in **Table S11**.

Primer efficiencies were calculated with standard curves generated by qPCR of serially diluted cDNA. ATF4 mRNA levels were quantified relative to RPL19 levels using the following equation:

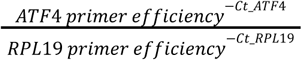

We similarly used this formula to calculate 18S rRNA values relative to RPL19 for the control comparison (**Figure S5**). Statistical analyses were carried out using paired t-tests, where paired samples correspond to RNA samples from Start-stop^WT^ and ΔStart-stop collected on the same day and qPCR reaction done in the same plate.

To test for the effect of the start-stop sequence in the human ATF4 5’ UTR on translation of uORF2 and uORF2 bypass under stress, we cloned the 5’ UTR sequence of human ATF4 from the transcription start site to the start codon of uORF2 into the dual-luciferase psiCHECK2 plasmid. This was done to ensure that uORF2 was in the same reading frame with the reporter gene hluc+ (firefly luciferase), allowing us to use firefly luciferase activity as a measure of translation initiation at the uORF2 start codon. uORF2 bypass under stress could now be measured as a decrease in translation of the uORF2-firefly fusion construct. We used renilla luciferase activity as a readout of global translation levels which we expected to be reduced under stress.

We also used the same 5’ UTR sequence, but mutated the AUG start codon of the start-stop to UUG to analyze the initiation activity at the uORF2 start codon in the absence of a start-stop. Lastly, we constructed reporter plasmids for the human ATF4 5’ UTR with uORF1 and uORF2 start codons mutated to AGG in a similar manner, with the ATF4 5’ UTR fused to the firefly reporter gene hluc+.

To determine bypass, we first normalized firefly luciferase and renilla luciferase activities by the transfection efficiency correction factors, and then calculated the change in normalized firefly in response to stress (i.e. bypass). We also corrected for the background change as measured by the normalized renilla luciferase activity.

## Results

### Start-stop elements efficiently stall ribosomes

We defined start-stops as sequence elements in the 5’ UTR of eukaryotic genes that consist of a start codon immediately followed by a stop codon, focusing only on canonical AUG start codons (**Figure 1A**). We defined control sequences as ‘AUUGGA’, ‘AUUGAA’, or ‘AUUGAG’, i.e., start-stop sequences with the inner two nucleotides swapped. Such start-stop and control sequences occur in the 5’ UTR of 1, 417 and 1, 837 human genes, respectively (**Table S1**). When examining a ribosome footprint dataset collected from HeLa cells providing data for ∼14, 000 human genes (26), we found that about two-thirds of the start-stop and control sequences were expressed at the mRNA level, and 312 and 122 of these had one or more ribosome protected fragment (RPF) associated with the element, respectively. To map reads, we required an RPF length of 28-30 nucleotides, and the read on a start-stop to be positioned with the start and stop codon localizing exactly to the P- and A-site, respectively (**Figure 1A**).

**Figure 1.**
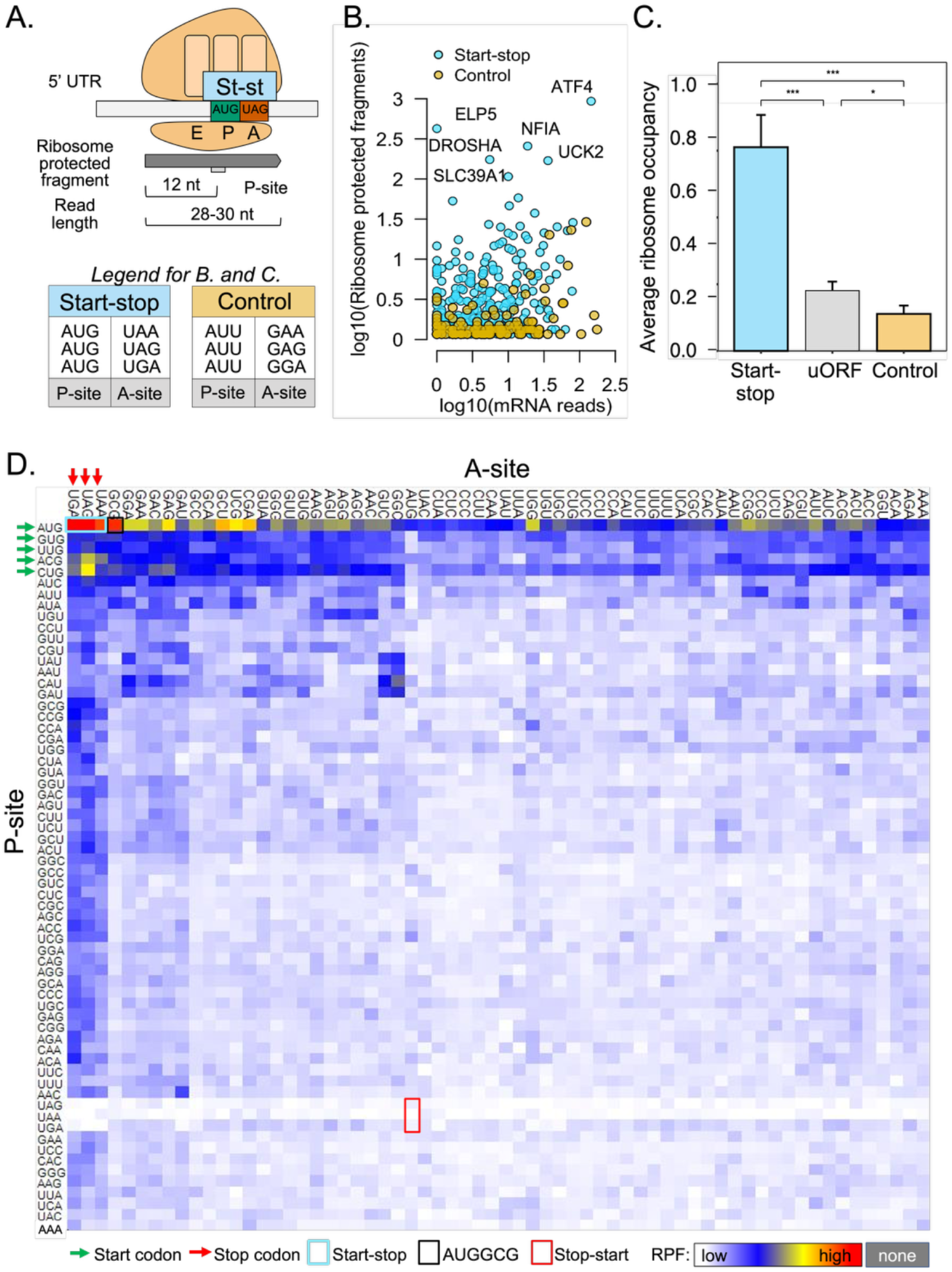
Start-stop elements are widespread and frequently occupied by a ribosome. **A.** The schematic illustrates the start-stop element in relationship to the bound ribosome. Ribosome footprints starting at the -12 position (P-site) have a typical length of 28-30 nucleotides. **B.** Scatter plot showing ribosome occupancy (RPF) vs. mRNA read depth for individual start-stops and control sequences in 5’ UTRs. **C.** Average ribosome read count on the AUG codons of start-stop and 9 nucleotide uORFs (any middle codon) and on the AUU of the control located to the 5’ UTRs. Data was taken from a ribosome footprint assay *without* the use of translation inhibitors (e.g. cycloheximide)(35). A t-test was performed to assess the statistical significance of the observed differences. Graphs show the mean and standard error of the mean. **Figure S1A** shows the extended data for thai plot. **D.** The heatmap dissects the 5’ UTR to display the fraction occupied for all P-site/A-site combinations (64 x 64 = 4, 096). Initiation P-sites, including non-canonical start codons, are highlighted with green arrows, while termination A-sites are denoted with red arrows. The AUGGCG hexamer is represented by the black box. Underlying data are shown in **Table S2. Figure S1B** shows related data, i.e. codon frequencies for the second codon in 9 nucleotide uORF. nt - nucleotides; * - *P-value* < 0.05; ** - *P-value* < 0.01; *** - *P-value* < 0.001

We compared ribosome coverage of start-stop elements to that of control sequences and found that start-stop containing sequences were substantially more often occupied by ribosomes than control sequences (**Figure 1B**), suggesting that ribosomes preferentially occupy start-stop sequences rather than other regions in the 5’UTR. The start-stop in *ATF4* showed the highest occupancy, followed by the start-stops in *UCK2*, *ELP5*, *SLC39A1*, *NFIA*, and *DROSHA*. This higher occupancy of start-stop elements held true for datasets collected without the use of translation inhibitors, e.g. cycloheximide (**Figure 1C**). Additionally, start-stop elements were significantly more often occupied than minimal uORFs, i.e. uORFs with only one additional codon (p-value < 0.001, **Figure 1C, Figure S1A**).

Next, we asked how ribosomes occupying a start-stop compared against ribosomes on other hexamers in the 5’ UTR. To do so, we calculated the frequency of ribosome occupancy for all P- and A-site codon combinations (N=4, 096) across all 28-30 nt footprints observed in the 5’ UTRs (**Figure 1D, Table S2**). In general, AUG codons were enriched in RPFs in P-site positions, and all three stop codons were enriched in the A-site. In contrast, ribosomes were depleted from combinations with stop codons in the P-site and AUG in the A-site (**Figure 1D**, “stop-start”, red box).

Notably, we observed the overall highest ribosome occupancy for the start-stop sequences, i.e., a combination of an AUG in the P-site and a stop codon in the A-site (**Figure 1D**, top left corner). In addition, one hexamer, AUGGCG, exhibited occupancy similar to that of start-stop sequences (**Figure 1D**, black box). This hexamer matches the favored translation initiation context in mammals (45, 46) and most likely represents the initiation site of uORFs in the 5’ UTR. Indeed, GCG was amongst the most frequent middle codons in minimal uORFs (**Figure S1B**). Further, we found high ribosome occupancy for near cognate start codons (GUG, UUG, ACG, and CUG, **Figure 1D**, green arrows) when positioned to the P-site of the hexamer. This effect was stronger with a stop codon positioned to the A-site, specifically for CUGUAG, CUGUGA, ACGUGA, ACGUAA, and ACGUAG. This observation suggested that these hexamers, consisting of near cognate start codons followed by a stop codon, might represent non-canonical start-stop elements.

### Start-stop elements repress translation of many signaling genes

To examine the properties of start-stop elements, we constructed a high-confidence core dataset in which start-stop and control elements were required to have >3 RPFs in two independent datasets (HeLa and HEK293) to qualify as ‘occupied’, or to have 0 RPFs in both datasets, but an expressed transcript with >5 RNA reads to qualify as ‘unoccupied’ (**Figure 2A**). Occupied start-stop sequences were significantly more abundant than occupied control sequences (occupied start-stop n = 88, occupied control sequence n = 7, unoccupied start-stop n = 21, unoccupied control sequence n = 85; p-value < 0.0001; **Figure 2A, Table S3**). We proceeded to analyze the properties of this core dataset (**Table S4**), and extrapolate to the extended dataset, as specified below.

**Figure 2.**
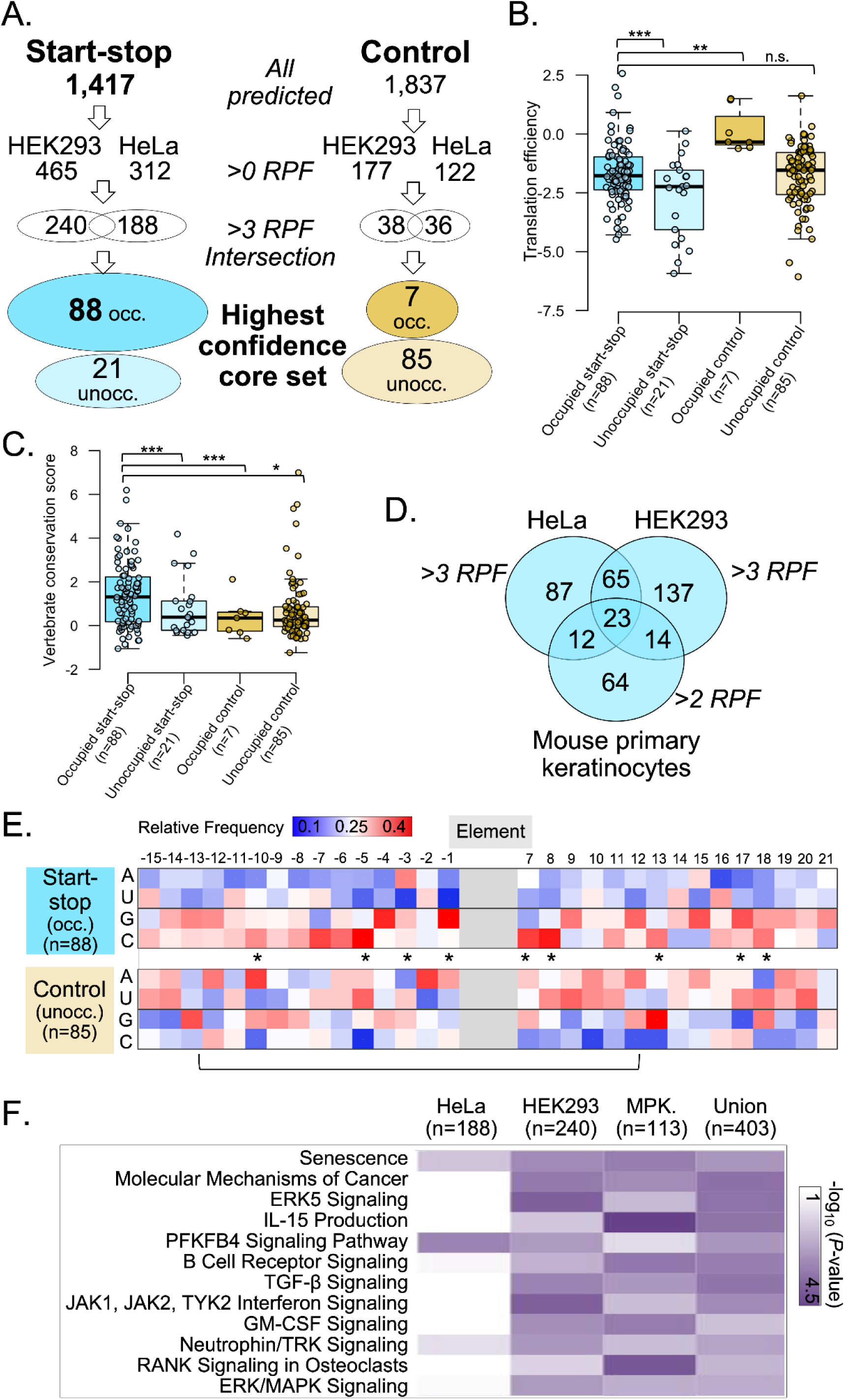
Start-stop elements are conserved and suppress downstream translation of many genes with key functions. **A.** Flow chart showing the number of genes with start-stop (stst) or control sequences predicted across the entire human genome (‘All predicted’), with at least one ribosome footprint (RPF) and more than 3 RPF in two HEK293 and HeLa datasets. A final ‘High confidence core set’ is defined by more than 3 RPF in *both* HEK293 and HeLa cells (‘occ.’ = occupied) or 0 RPF, but >5 RNA reads in *both* datasets (‘unocc.’ = unoccupied). All predicted start-stop and control genes are listed in **Table S1**. The high-confidence core set is defined and described in **Tables S3/S4**. * - *P-value* < 0.05; ** - *P-value* < 0.01; *** - *P-value* < 0.001; n.s. - not significant **B.** Translation efficiencies of genes containing a core start-stop or control sequences. Translation efficiency of CDS was calculated from ribosome profiling and RNA-seq data in HeLa cells (26). Related data are shown in **Figure S2A**: translation efficiencies of start-stop and control genes. **C.** Sequence conservation in vertebrates compared among core start-stops and controls, based on occupancy with ribosomes. Phylop-based conservation scorers were obtained using 100 vertebrate species, downloaded from the UCSC browser (https://genome.ucsc.edu). **Tables S3/S4** show the genes and properties. A *Drosophila* example of a start-stop containing gene is shown in **Figure S2B**. **D.** Venn diagram showing overlap of start-stop-containing genes in HeLa (26), HEK293 (30), and mouse primary keratinocytes (33). Human and mouse data were filtered for start-stop sequences with at least three and two RPF, respectively. **Tables S5** shows the genes. **E.** Relative frequencies of each nucleotide at positions ± 15 upstream and downstream of start-stops and controls. A * marks position with significant differences in nucleotide frequencies (chi-square test, p-value < 0.05). The bracket marks the approximate region covered by an 80S ribosome. Nucleotide frequencies for the four high-confidence core datasets are shown in **Figure S2C. Tables S3/S4** show the genes and properties. **F.** Heatmap displaying enriched pathways for each cell type and the union, using start-stop sequences with at least three and two RPF in human and mouse, respectively, as shown in Figure 2D and **Table S5**.

Based on the high ribosome occupancy of start-stop elements, we hypothesized that start-stops reduce ribosome procession and therefore lower translation of the downstream open reading frame (ORF). To test this hypothesis, we examined the translation efficiency of the main ORF, as estimated from the ribosome footprint data. Indeed, we found that translation efficiency of the genes with occupied start-stops was significantly lower than that of genes with occupied control sequences in the core dataset (p-value < 0.001, **Figure 2B**). This observation held true also when examining the expanded dataset with more genes (**Figure S2A**), suggesting that the presence of a start-stop in the 5’UTR indeed represses translation of the downstream ORF. This analysis also revealed that mRNAs with unoccupied start-stops had a lower translation efficiency on average (**Figure 2B**). We speculate that these may be cases where the start-stop acts as a fail-safe in 5’ UTR: the start-stop functions in addition to other elements that repress translation without being occupied itself.

Next, we hypothesized that if start-stop sequences played important roles in translation regulation, their sequence should be evolutionarily conserved more than that of the control sequences which might be non-functional. We found that indeed the sequence of the core set of occupied start-stop hexamers was more conserved across 100 vertebrate species than that of the controls, supporting their function as a regulatory element (p-value < 0.01, **Figure 2C**). This conservation is remarkable given the rapid evolution of untranslated regions (47).

The sequence conservation was confirmed by the significant overlap of human start-stop containing genes with those from mouse (p-value < 0.01, **Figure 2D**). When examining the ribo-seq dataset collected from mouse primary keratinocytes (33), we identified 113 genes with a ribosome positioned to the start-stop with >2 RPF counts (**Table S5**). Note that the cutoff for mouse was less stringent than that for the human datasets (>3 RPF) as the overall coverage was lower for the mouse data. Of these occupied mouse start-stop genes, 23 overlapped with the core set of occupied human start-stop genes which represented the intersection between the HeLa and HEK293 data (**Figure 2D**). This overlap is substantial given the stringent cutoffs that we used, the different cell lines considered, and the different experimental conditions. Finally, we found many start-stop-containing genes in the single cell eukaryote *S. cerevisiae* (**Table S6**), and a specific example in *D. melanogaster* discussed below (**Figure S2B**).

To further investigate the possible functionality of start-stop elements through halting the scanning 48S PIC and subsequent translation initiation, we examined the surrounding sequence. For start-stop sequences, it is impossible to achieve a ‘perfect’ translation initiation context, as the nucleotide at the +4 position is a U when any stop codon follows a start, while in the Kozak consensus sequence the +4 position is preferably a G or A (45). Nevertheless, we found that occupied core start-stops had a sequence context consistent with efficient initiation at other positions, with several statistically significant differences to the controls (p-value < 0.05, **Figure 2E**). For example, occupied start-stop elements from the core set preferred A or G at the -3 position, while unoccupied start-stops and unoccupied control sequences had mixed preferences at the -3 position (**Figure 2E, Figure S2C**). Occupied start-stops also strongly preferred G at the -1 position compared to all other groups. We also observed a general GC enrichment within +/- 15 nucleotides surrounding the occupied start-stops, contrasting the relative AU rich regions surrounding unoccupied control sequences (**Figure 2E**). The GC enrichment could indicate RNA secondary structures (48) that slow ribosomes as they scan to reinforce initiation immediately upstream of their occurrence in the sequence, or potentially mediate interactions with additional translation factors.

We manually examined the core set of occupied control sequences for potential reasons for their ribosome occupancy (**Table S7**). Two control sequences demonstrating occupancy were located within a longer uORF - which would explain the occupancy with ribosomes. In the other occupied control sequences, the AUU in the element could potentially serve as a non-canonical initiation site (49), providing another explanation for ribosome occupancy. By definition, all control sequences had a G in the +4 position, in support of the Kozak sequence (**Table S7**). All control sequences also had A, G, or C at the -1 position, and 5 had A or G at the -3 position, further strengthening the Kozak context for possible initiation at the AUU. However, despite these explanations for ribosome occupancy of the control sequences, their low abundance (n = 7) compared to that of occupied start-stops (n = 88) illustrates the strong preference of ribosomes to start-stop elements.

Finally, start-stop-containing genes were significantly biased in their functions, both within the high-confidence dataset and the expanded set (**Figure 2F**, **Tables S8, S9**). They participated in various signaling pathways, e.g. ERK5, TGF-β, and PFKFB4, while being implicated in cancer (p-value < 0.001). Start-stop containing genes were also significantly enriched in transcription factors (p-value < 0.001), such as *ATF4*, *NFIA*, *JARID2*, and *SMAD6/7*. *ATF4* is discussed in more detail below. NFIA is an important transcription factor involved in gliogenesis with some links to the stress response (50, 51). *JARID2* is a transcription repressor with roles in development (52). Smad proteins play a key role in TGF-β signaling regulation (53). The start-stop sequence was conserved in one transcript isoform of the *Drosophila melanogaster* homologous gene Mothers against dpp (Mad). When expressed in S2 cells, the start-stop-containing isoform is occupied by ribosomes (54) **(Figure S2B)** supporting the likely functionality of the element and suggesting that expression of transcript variants with and without the start-stop element in diseased tissue might contribute to the dysregulation of SMAD7 protein levels frequently observed in cancers (54).

### Start-stop elements are often part of complex regulatory scenarios

Next, we examined typical start-stop gene architectures to investigate their possible modus operandi (**Figure 3A**). We found start-stops occur in a variety of transcript-specific contexts. For instance, in some genes such as *MORF4L1, UCK2, MEN1, NFIA, CACUL1,* and *CTXN1*, the start-stop appeared to be the only detectable sequence element in the 5’ UTR. Many other genes with start-stops had more complex architectures that included additional uORFs, e.g. *ATF4, GPR37, DROSHA, ELP5, IFNAR1, SLC39A1,* and *PSPC1* (**Figure 3A**). In these genes, the regulatory mode is likely complex. In some genes, e.g. *ATF4* and *ELP5*, the start-stop occurred close to the 5’cap (**Figure 3A**), potentially obstructing scanning by the small subunit. In other genes, e.g. *DROSHA* and *IFNAR1*, the start-stop was located in close proximity to the CDS, likely both reducing ribosome scanning and repressing reinitiation downstream of the start-stop by impairing the ribosome’s ability to reinitiate fast enough for efficient translation. This effect is due to an insufficient time/distance for the post-termination ribosome to reacquire the ternary complex (TC) whose concentration is frequently limiting.

**Figure 3.**
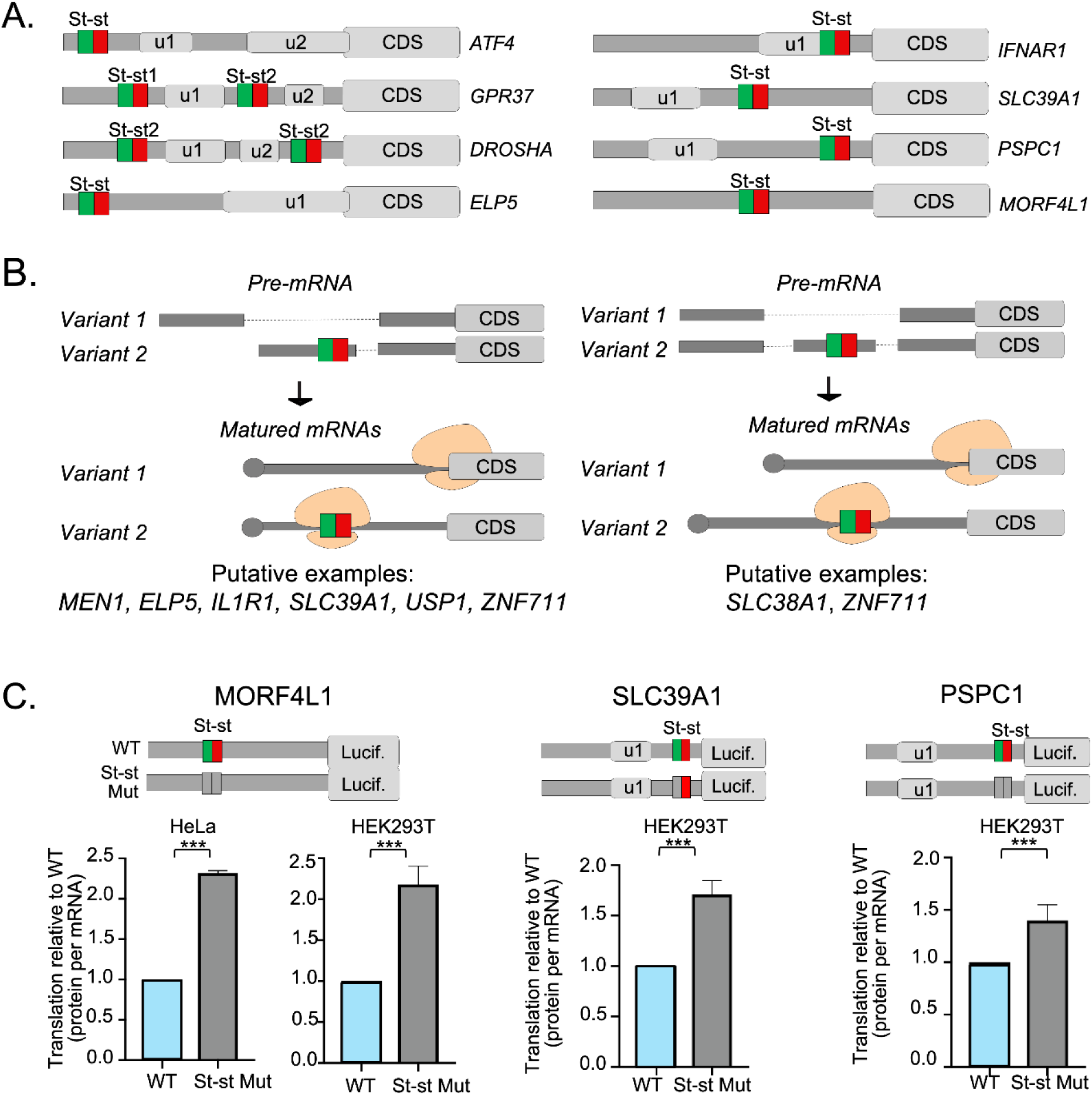
Start-stop elements occur in diverse 5’ UTR architectures. **A.** Diagrams showing the positioning of start-stops within 5’ UTRs relative to uORFs for sample genes, not drawn to scale. **B.** Diagrams showing example genes with highly occupied start-stop sequences that only occur in some transcript variants, but not in others. **Figure S3A/S3B** show example genes with their transcript expression data. **C.** Schematics of start-stop containing genes tested for the effect of the start-stop on translation of the CDS. Relative luciferase activity was measured in HEK293T transfected with plasmids expressing luciferase fused to the gene’s 5’UTR. Experiments included Hela cells for MORF4L1. All experiments were conducted in triplicate and show the translation efficiency, i.e. the measured luciferase activity normalized by the mRNA expression level. Measurements were also normalized to the wild-type (intact start-stop). All graphs show means and standard errors of the mean. We used t-tests to assess significance. **Figure S3C** shows the separate protein and RNA measurements. * - *P-value* < 0.05; ** - *P-value* < 0.01; *** - *P-value* < 0.001. CDS - coding region; Lucif. - luciferase reporter; n.s. - not significant; WT - wild-type

Further, we examined instances of start-stop genes for the positioning of the element with respect to the exon-intron structure of the 5’ UTR. We identified several cases in which either alternative transcription start sites or alternative splicing produces transcript variants for the same gene that either included or excluded the start-stop (**Figure 3B**). For example, in several genes the start-stop was located in the first exon, but alternative transcript variants used different first exons (**Figure 3B**, left). Examples of these genes included *MEN1, ELP5, IL1R1, SLC39A1,* and *USP1*. We observed high coverage of the *MEN1* start-stop in both human and mouse (**Table S5**). *ELP5* and *SLC39A1* were highly occupied in HeLa and covered in HEK293 (**Table S5**); the mouse orthologs had no start-stop, but short uORFs instead. The start-stops in *IL1R1* and *USP1* were highly occupied in HeLa, but less so in the other datasets. In other examples, e.g. *SLC38A1* and *ZNF711*, the start-stop was located in an internal exon in the 5’ UTR, and the exon was either present or absent in alternative variants (**Figure 3B**, right). In both genes, the start-stop showed high ribosome occupancy in the human datasets.

We hypothesized that transcript variants including the start-stop would be translationally repressed, while transcript variants excluding the start-stop would be translated efficiently - providing a potential mechanism of expression regulation at the level of translation. Indeed, we manually identified cases in which this might be taking place, i.e. genes where different transcript variants that include or exclude the start-stop element and where the transcript variants show tissue specific expression patterns. One example was USP1 shown in **Figure S3A**. USP1 is a deubiquitination enzyme with diverse roles, e.g. in the regulation of DNA damage repair (55). It had four transcript variants with several translation regulatory elements, but only one contains the start-stop bearing exon. As USP1 protein is overabundant in cancer (55), dysregulation of its expression might be, in part, due to switching to transcript variants without the translation repressive start-stop.

Another example arose from ZNF711, which is of unknown function but with similarity to transcription factors. It is linked to brain development and its mutation causes intellectual disability (56, 57). ZNF711 had a complex 5’ UTR with several exons and three start-stop elements, creating five distinct transcript variants with three different combinations of start-stop inclusion or exclusion (**Figure S3B**). All five transcript variants were expressed with distinct tissue-specific patterns.

Next, we demonstrated the impact of the start-stop on translation of several example genes, i.e. *MORF4L1, SLC39A1,* and *PSPC1*. Mutating the start-stop significantly de-repressed translation in each case (p-value < 0.001, **Figure 3C, Figure S3C**), supporting our hypothesis that start-stops impede ribosomes as they scan through the 5’ UTR, while the mutation removes this blockade. (Note that graphs from the luciferase assays always show the protein levels normalized to the levels of corresponding mRNAs, unless specified otherwise.) *MORF4L1* is a little-characterized gene involved in cancer cell migration (58), and conserved in mouse and human. Mutating its start-stop significantly increased translation in both HeLa and HEK293T cells (p-value < 0.001). Mutating the start-stop in SLC939A1 and PSPC1 increased translation of the reporter ∼1.5-fold, suggesting that the start-stop is functional (p-value < 0.001, **Figure 3C, Figure S3A, S3B**). SLC39A1 is a member of the zinc-iron permease family and linked to several cancers (59, 60). Its 5’ UTR variants can include or exclude the start-stop (**Figure 3B**), while encoding the same protein (61). A large-scale screen identified an internal ribosome entry site also present in SLC39A1’s 5’ UTR creating a more complex regulatory model which might explain the comparatively low impact of the start-stop (62). Paraspeckle component protein 1 (PSPC1) marks mammal-specific RNA–protein nuclear bodies that regulate gene expression. The protein is linked to several cancers (63).

The mutated start-stop had no effect on transcript abundance for *MORF4L1*, while it slightly lowered transcript abundance for *SLC39A1* and *PSCP1*, suggesting that the start-stop might stabilize the mRNA (**Figure S3C**). This finding was surprising given that canonical (short) uORFs have been shown to destabilize the mRNA in reporter constructs (64).

We generalized the analysis of start-stop positioning across the entire dataset and found that occupied start-stop elements tended to occur closer to the 5’ cap than unoccupied control sequences (not shown). However, occupied control sequences and occupied minimal uORFs were also closer to the 5’ cap than unoccupied sequences, therefore suggesting that cap proximity is a general property of functional translation regulatory elements capturing the first scanning ribosomes and not a start-stop specific property.

### Start-stop elements repress translation more so than uORFs

Start-stops are related to canonical uORFs, but function without an elongation step. We hypothesized that this could take place through two different scenarios. Upon initiation, the ribosome must immediately enter termination, but is unable to complete termination perhaps due to the lack of the P-site tRNA-polypeptide hydrolysis step (Model (I), **Figure 4A**). Release factors would repeatedly associate and dissociate and the 80S ribosome would remain ‘stuck’ on the start-stop site.

**Figure 4.**
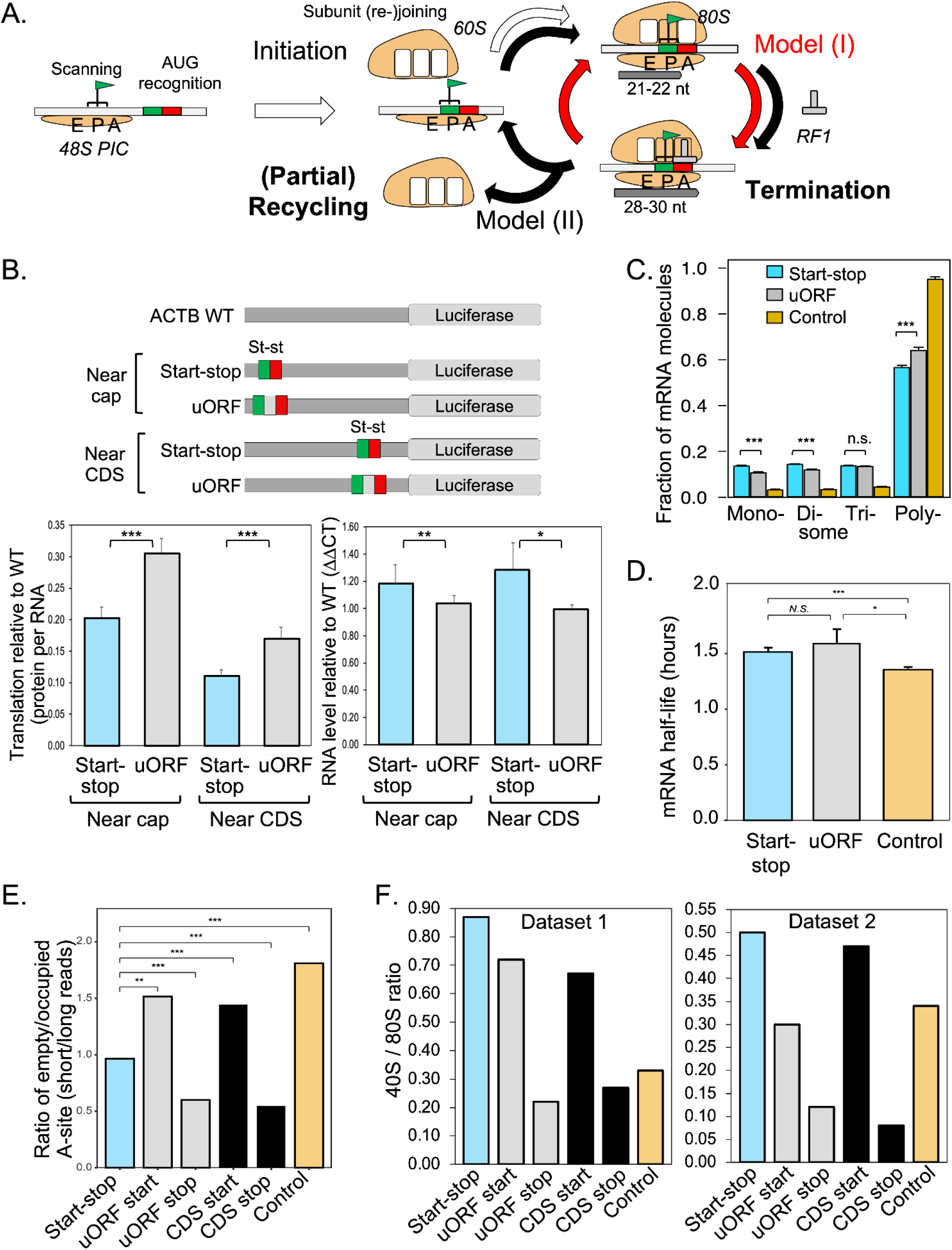
Start-stops are distinct from canonical uORFs. **A.** Diagram explaining models proposed for ribosomes stalling on start-stop elements. A scanning 48S PIC would initiate upon detecting the start-stop’s AUG and recruiting a 60S subunit (white arrow). However, ribosomes would not enter elongation, but immediately proceed towards termination by recruitment of termination factors. Termination might be incomplete and cause eRFs to come on and off without actually completing the termination phase (Model (I); red arrows). Alternatively, termination might complete all steps and proceed into partial recycling during which the 60S subunit detaches but the 40S subunit lingers at the start-stop site (Model (II); black arrows). The 60S subunit will then rejoin to enter a new cycle. Analysis of ribosome footprint width (read length) and 40S footprints compared to 80S footprints suggested that Model (II) is more likely. **B.** Gain-of-function constructs probing the effect of start-stop elements on translation and mRNA stability. The diagrams on the top explain the constructs used. We inserted either start-stop (‘Start-stop’) or 9 nucleotide uORF (‘uORF’, defined as AUGGCGUGA) sequences into the ACTB 5’ UTR used for the luciferase reporter. The unmodified ACTB 5’ UTR represents the wild-type which all measurements had been normalized to. The bar plot on the left shows the translation efficiency for the different constructs as calculated from the ratio of the protein and RNA level measured by luciferase assays and by RT-qPCR, respectively. All results are relative to the wild-type ACTB 5’UTR set to 1 (not shown). The bar plot on the right shows the mRNA levels as measured by RT-qPCR for the different constructs. All experiments were conducted in triplicate. We performed a t-test to determine the significance of the differences. **Figure S4A** shows related data with a 21 nucleotide uORF. **C.** Start-stop elements, minimal uORFs, and control sequences distributed across mono-, di-, and polysome fractions. The plot shows the fraction of RNA sequences observed with the ribosome reads at -12 position relative to the AUG of start-stop and 9 nucleotide uORF (any middle codon), and to the AUU of the control in the 5’ UTRs. The control sequences were the three possible start-stop elements with nucleotides 3 and 4 swapped, eliminating the start and stop codon. Data was taken from an unbiased screen of the differential ribosome binding to same-length RNA leaders of randomized sequence (39). The different sections mark the mono-, di-, tri-, and polysome fractions as per (‘mono’, ‘di’, ‘tri’, ‘poly’, respectively). We performed a t-test to determine the significance of the differences. **Figure S4B** shows a similar analysis performed on a different dataset. **D.** Start-stop elements, minimal uORFs, and control sequences plotted with their RNA half-life. Start-stops, uORFs, and control sequences were defined as described for **C.** The data were taken from (40) which screened a library of randomly designed 10 nucleotide sequences for their effect on transcript stability. **E.** Ratio of short (21-22 nucleotides) and long (28-30 nucleotides) ribosome footprint counts signifying empty and occupied A-sites, respectively, across start-stop elements, uORFs, CDSs, and control sequences. Footprint data was taken from a dataset produced without the use of cycloheximide (35). Ribosome reads were counted if they were located to the -12 position relative to the AUG of the start-stop, the 9 nucleotide uORF (any middle codon), and the CDS or to the -15 positions relative to the stop codon of the uORF and of the CDS, respectively, and at the -12 position relative to AUU of the control sequence. We performed Fisher’s exact test to determine the significance of the differences. **Figure S4C** shows the deconvoluted data. **F.** Ratio of 40S/80S ribosome footprint counts taken from Dataset 1 (12) and 2 (21). Footprints were counted when positioned to the respective elements described in **E.** Due to low 40S read counts, data was aggregated for the respective sections prior to calculating the ratio. Plots in **B.-D.** show means and error of the mean. CDS - coding region; * - *P-value* < 0.05; ** - *P-value* < 0.01; *** - *P-value* < 0.001.

Alternatively, the ribosome initiates and enters termination, completes it and undergoes the first step of recycling, which involves detachment of the 60S subunit (Model (II), **Figure 4A**). The 60S subunit would then rejoin the 40S subunit that still has the Met-tRNA_i_^Met^ in the P-site, therefore building an elongation-capable, fully assembled ribosome. However, since the ribosome cannot enter elongation, as it is already positioned with the stop codon in the A-site, this cycle of termination and partial recycling would be repeated several times until the 80S ribosome resumes scanning or fully dissociates.

We probed these models by examining properties of start-stops in comparison to canonical open reading frames. We first investigated the translation effect of start-stop elements in comparison to minimal uORFs. As properties of uORFs vary depending on their length and codon composition (64), we focused on the shortest possible (‘minimal’) uORF, i.e. consisting of 9 nucleotides. For targeted experiments, we chose the uORF’s middle codon to be GCG as it had one of the highest average ribosome occupancies amongst all ‘middle’ codons in human 9-nucleotide uORFs (**Figure 1C, Figure S1B**), consistent with a strong Kozak sequence. High initiation at a uORF represses translation at the downstream ORF, underlying the known translation inhibitory role of many uORFs (64). We argue that in this way we did not bias the uORF towards suboptimal initiation such as found in start-stops and therefore a reduced impact on the downstream translation.

We conducted a series of gain-of-function experiments in which we inserted start-stop elements as well as the minimal uORF into the 5’ UTR sequence of *ACTB* (**Figure 4B**). *ACTB’s* 5’ UTR has a length of 84 nucleotides and does not - to the best of our knowledge - contain any known translation regulatory elements. To mimic scenarios we had observed in individual start-stop-containing genes (**Figure 3A**), we inserted the start-stop or minimal uORF either near the 5’ cap or the coding region, i.e. 77 and 30 nucleotides upstream of the CDS, respectively (**Figure 4B**). Using quantitative RT-PCR and luciferase reporters, we estimated the impact of the elements on transcript and protein expression levels, respectively.

Indeed, both the start-stop and the uORF repressed translation of the reporter compared to the wild-type ACTB 5’ UTR (**Figure 4B**). Translation is measured as the change in protein levels normalized by the RNA level. This repression could be attributed to the elements hindering the steady ribosome ‘flux’ (scanning) along the 5’UTR towards the CDS. Further, for both the start-stop and the uORF, the effect was about two times stronger when the elements were positioned close to the CDS (**Figure 4B**), suggesting that not only was ribosome scanning (‘flux’) affected, but reinitiation was decreased if the downstream AUG was near the end of the respective element, as expected based on insufficient time to recruit a new TC (10, 65). Comparing these two different placements, the start-stop and uORF seemed to affect downstream re-initiation at the CDS to a similar extent.

The effect on observed translation (protein levels normalized by mRNA levels) was not due to changes in mRNA stability, even though start-stops and uORFs differed in their effect on mRNA levels (**Figure 4B**). The *ACTB* 5’ UTR transcript with an inserted minimal uORF remained at wild-type levels. In comparison, inserting a start-stop element slightly, although not significantly, stabilized the mRNA (∼1.3-fold). The finding was consistent with what we had observed for *SLC39A1* and *PSPC1*, where mutating the start-stop resulted in decreased transcript levels (**Figure S3A**).

Notably, we observed significantly stronger translation repression when inserting a start-stop compared to a uORF, regardless of the position within the 5’ UTR (p-value < 0.001, **Figure 4B**). This effect held true even for a longer uORF (**Figure S4A**). Inserting a start-stop led to a 5- to 10-fold reduction in translation, while we observed only a 3- to 6-fold reduction for inserted minimal uORFs. This observation was remarkable as the Kozak sequence for the start-stop was less optimal than that for the uORF as discussed above, due to the U at the +4 position in the start-stop. It suggested that the start-stops reduced ribosome ‘flux’ more so than strong-initiator minimal uORFs.

Next, we tested for the same characteristics in data from large-scale studies. **Figure 4C** shows the analysis of data from a screen measuring the impact of different 5’ UTR sequences on translation efficiency, as estimated from the distribution of the reporter sequences across different fractions of a polysome profile (39). The sequences comprised randomized stretches fused to a shared 5’ UTR. We examined these randomized sequence stretches for the presence of start-stops, minimal uORFs, or control sequences. Note that for this analysis, we considered 9-nucleotide uORFs with any middle codon, with the exception of an AUG (i.e. AUGAUG-stop).

Similar to the results from the gain-of-function experiments (**Figure 4B**), both start-stops and minimal uORFs repressed translation compared to the control, i.e. shifting sequences to the monosome fraction (**Figure 4C**). However, start-stops had a significantly stronger effect than minimal uORFs (p-value < 0.001). We confirmed these findings with data from another large-scale screen of randomized sequences, demonstrating a similar trend: start-stops were present in monosomes significantly more often than uORFs (p-value < 0.01, **Figure S4B**). Further, sequences with either a start-stop or a minimal uORF had slightly higher mRNA levels than control sequences (p-value < 0.05, **Figure 4D**), consistent with the above finding that start-stop elements did not seem to trigger higher RNA turnover.

### Start-stop elements might stall ribosomes through a non-canonical mechanism

Next, we tested start-stops for characteristics that could suggest a mechanism of the ribosome stalling that we observed (**Figure 1B, 1C**). As such, we first tested if ribosomes positioned onto start-stops showed biases with respect to A-site occupancy. An empty A-site is consistent with an initiating ribosome and with an elongating ribosome post translocation; an occupied A-site is consistent with a terminating ribosome and with an elongating ribosome post accommodation and peptide bond formation (**Figure 4A**). To test for any biases, we exploited the fact that the ribosome footprint length is representative of A-site occupancy: ribosomes with empty A-sites produced shorter reads (21-22 nucleotides) than ribosomes with occupied A-sites (28-30 nucleotides)(35). The data used had been acquired for HeLa cells without any inhibitors (35).

In this dataset, ribosomes on start codons of minimal uORFs or main open reading frames (CDS) had empty A-sites more often than ribosomes on stop codons (p-value < 0.01, **Figure 4E, Figure S4C**), indicating that ribosomes waiting at start codons for the first elongation ternary complex composed of eEF1A, GTP and an aminoacylated tRNA (i.e. the codon sampling and accommodation steps of elongation) were more common than ribosomes waiting for release factors at the stop codon. In other words it seems that the conversion rate of the just assembled 80S initiation complex into an elongation competent 80S ribosome was lower than that of the fully engaged elongation complex into a termination complex. In comparison, ribosomes stalled at the start-stop elements had read length distributions significantly different from any of the other distributions (**Figure S4C**) with summary values between those observed for the start and stop codon at uORFs/CDSs (p-value < 0.001, **Figure 4E**). This finding suggested that ribosomes on start-stops did not have any obvious bias towards stages with occupied or empty A-sites.

Next, we tested if start-stops had a bias in 40S subunits (or 48S PIC) compared to a fully assembled 80S ribosome bound to the site. A 40S footprint could indicate a scanning PIC, or a partially recycled ribosome after termination (**Figure 4A**). We analyzed two datasets which collected footprints of both the 40S and 80S ribosome for mammalian cells (12, 21) and compared the 40S/80S read ratio between start-stop elements, the control, and the start and stop codon of minimal uORFs and the CDSs. As 40S footprint data was comparatively sparse, we pooled read information in a metagene analysis prior to calculating the 40S/80S ratio, rather than calculating ratios for individual elements.

In both datasets, start-stops had a higher 40S/80S ratio than any other group (**Figure 4F),** suggesting that they were comparatively frequently occupied by the small subunit only. The two datasets produced different absolute 40S/80S ratios due to differences in the underlying methods. However, within each dataset, stop-codons at uORFs and the CDS had a comparatively low 40S/80S ratio, consistent with a small amount of post-termination ‘lingering’ of the 40S subunit. In comparison, start-codons at uORFs or coding regions showed a relatively high 40S/80S ratio consistent with a PIC waiting at the start codon to recruit the 60S subunit and initiate translation. Surprisingly, the 40S fraction at start-stops was even higher in both datasets (**Figure 4F**), even if the difference was small for Dataset 2 (21). This finding, combined with the above results, suggested that the ribosome on start-stops underwent cycles of termination and partial ribosomal recycling including temporary detachment of the 60S subunit (**Figure 4A**).

### A start-stop element in the human ATF4 gene supports the uORF-driven delayed reinitiation

Finally, we examined the role of the start-stop element in *ATF4* in more detail, complementing related work we have published recently (23). *ATF4* is the quintessential example of uORF-mediated translation induction in metazoans (4, 10, 65): two sequential uORFs repress translation under normal conditions and induce translation during the integrated stress response, through a mechanism of delayed reinitiation combined with stem-looped-induced ribosome queuing (23). In this mechanism, the preinitiation complex scans the 5’ UTR until it recognizes the start codon of the first uORF (uORF1), and the short ORF is translated. Following termination at this uORF, the small ribosomal subunit remains associated with the mRNA and continues scanning downstream. Downstream reinitiation is dependent on ternary complex abundance: when TC levels are high, i.e. under normal conditions, TC recruitment occurs rapidly and reinitiation is favored at the next actionable start codon, which corresponds to the second uORF (uORF2). This second uORF overlaps with the main ORF. Therefore, termination of uORF2 occurs downstream of the main *ATF4* start codon, suppressing ATF4 protein synthesis. During stress conditions, when TC abundance is lower, reinitiation rates at the uORF2are reduced, promoting bypass of its start codon. However, as uORF2 is very long (59 amino acids in total), the small subunit eventually recruits a TC and the scanning ribosomes can reinitiate further downstream at the main ORF, thereby inducing ATF4 translation (7, 8). While this model has been well-established, several observations contrast the original model’s predictions (26, 66, 67) and have triggered additional investigation that revealed an additional regulatory layer in a form of stem-loop-induced ribosome queuing in the uORF2/ATF4 overlap region (23).

The human *ATF4* gene contains a start-stop element located upstream of uORF1, with very high ribosome occupancy in the HeLa data under both normal and stress conditions (**Figure 5A**)(12, 26). While the start-stop sequence is highly conserved across vertebrates (**Figure S5A**), it does not occur in the mouse Atf4 sequence where the original delayed reinitiation model had been discovered (7, 8). Prior work examined the start-stop in the human *ATF4* with respect to factors that impact reinitiation *downstream* of the element (21, 68, 69). Specifically, it showed that reinitiation after the start-stop depended on the correct function and sufficient levels of the reinitiation factors eIF2D and DENR/MCTS1 (21, 68, 69). These studies proposed a mechanism in which DENR/MCTS1 and/or eIF2D are required for release of the deacylated tRNA from the ribosome to enable recycling of the large subunit, continued scanning of the small subunit, and downstream reinitiation (21).

**Figure 5.**
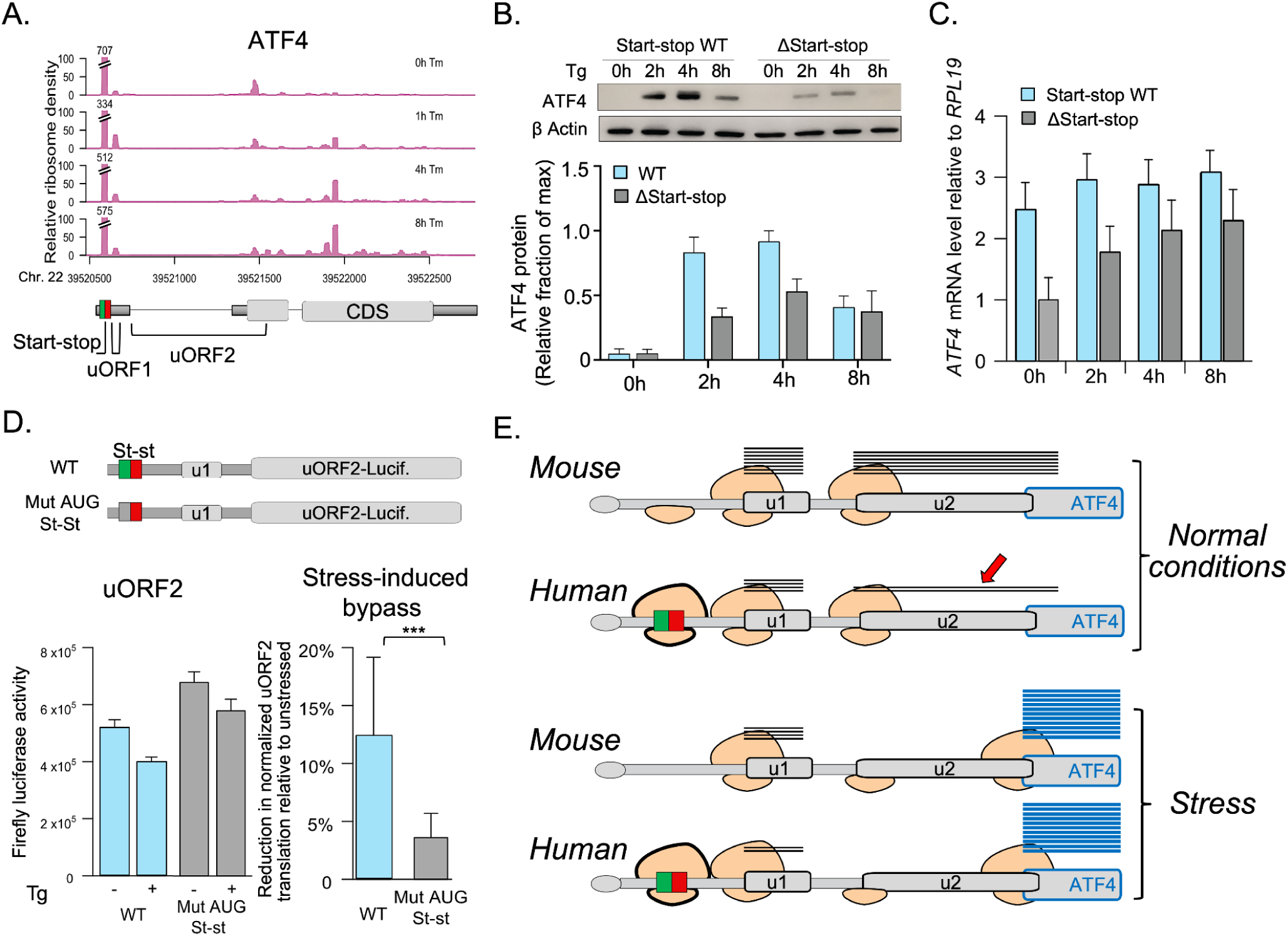
The start-stop in the human ATF4 gene supports the uORF-driven model of translation control. **A.** Relative ribosome density displayed as a function of position along the ATF4 gene. Profiles correspond to 0, 1, 4, and 8 hours post treatment with the stressor tunicamycin (Tm) in HeLa cells (26). The coordinates (GRCh38/hg38) along chromosome 22 are labeled; gene schematic indicates UTR (grey boxes), CDS (grey boxes), and introns (black lines). The positions of the start-stop, uORFs, and CDS are annotated in the diagram below. **Figures S5A** and **S5B** show the sequence conservation of ATF4’s start-stop and ribosome footprints on the start-stop across different datasets, respectively. **B.** Representative Western blots against ATF4 and loading control from CRISPR-edited HAP1 cells before and after treatment with thapsigargin (Tg, 1uM). Graphs below show the ATF4 protein levels normalized to the loading control (% of max) obtained from three replicate measurements. **Figure S5C** shows related data for the stop-stop mutation of the start-stop. **C.** ATF4 mRNA levels measured by qPCR in three replicate measurements, corresponding to the samples from panel **B**. **D.** Diagrams and data measuring uORF2 translation in a uORF2-luciferase fusion construct. The graph on the left shows the firefly luciferase activity of the two constructs; the plot on the right shows the change in uORF2 bypass after Tg treatment. Values are firefly values of the fusion construct normalized the renilla luciferase measurements. **Figures S5D-S5G** show additional constructs and datatypes as described in the text. **E.** Diagram for the mouse and human ATF4 gene without and with a start-stop element (stst) upstream of the uORFs (u1, u2), respectively. Positions and sizes are not drawn to scale. The grey and blue lines represent the peptides produced from the uORFs and the CDS, respectively. The red arrow points to the amount of peptide produced from uORF2, which is much lower in human than in mouse, thanks to the start-stop reducing overall ribosome load. **Figures S5H/S5I** show the simple mathematical model illustrating the start-stop function in the human *ATF4*. Extended data are shown in **Figure S5.** Graphs in **B.-D.** show means and standard error of the mean. CDS - coding region. * - *P-value* < 0.1; ** - *P-value* < 0.01; *** - *P-value* < 0.001.

While these findings explain how translation can resume after the start-stop in the human *ATF4* gene, they did not explain the specific impact of the start-stop on the ATF4 translation system itself. Given that the ribosome read counts at the start-stop are higher than anywhere else in the 5’ UTR (**Figure 5A**), independent of the use of inhibitors (**Figure S5B**), we hypothesized that the *ATF4*’s start-stop has a significant impact on the overall ribosome flow and translation of both the upstream and main open reading frames. Specifically, we hypothesized that the start-stop would support the delayed reinitiation model, i.e. ensure translation suppression of the main open reading frame under normal conditions, but reduce wasteful production of the long peptide encoded by the second uORF and therefore improve cellular economy. To test this hypothesis, we examined a series of luciferase reporter constructs, as well as CRISPR-Cas9 edited Hap1 cell lines.

Indeed, removing the start-stop by CRISPR-Cas9 (6 nucleotide deletion) resulted in little observable changes in basal ATF4 levels as measured by Western blotting (**Figure 5B**), consistent with the established model in which the two uORFs are sufficient to repress ATF4 translation under normal conditions (23). The results were the same when modifying the start-stop to a stop-stop (UAGUAG) sequence (**Figure S5C**). The result was also confirmed by a reporter construct which contained the start-stop but no other functional uORF (**Figure S5D**): the start-stop reduced the overall ribosome flow and hence translation of downstream elements.

Further, similar to what we observed for the start-stops in general (**Figure 4C, Figure S2**), mutating the start-stop in *ATF4* resulted in lower RNA levels compared to the wild-type (**Figure 5C**, **Figure S5E-F**). This result also aligned with previous work that had shown the translation of second uORF triggers *ATF4* mRNA degradation via nonsense-mediated mRNA decay (NMD)(70). Thus, reducing translation of the second uORF through ribosomes stalling at the upstream start-stop under normal conditions would lower NMD-related degradation and consequently increase *ATF4* mRNA half-lives - as we observed.

Unexpectedly, we observed a second role of the start-stop: mutating it resulted in reduced ATF4 induction under stress (**Figure 5B**), independent of the changes in transcript levels (**Figure 5C**). We hypothesized that the reduced inducibility in our system was due to the start-stop supporting stress-induced bypass of the second uORF. To test this hypothesis, we designed reporters that fused the human *ATF4* 5’ UTR up to the second uORF’s start codon to the firefly luciferase coding sequence (**Figure 5D**). The reporter also included a renilla luciferase expressed from the same plasmid as firefly luciferase. The renilla luciferase intensity served as an internal control to monitor overall translation levels under normal conditions and under stress (**Figure S5G**).

This construct allowed us to quantify translation reinitiation at uORF2 (through the firefly readout). We hypothesized that the start-stop would not affect reinitiation rates *after* uORF1. However, the start-stop as a road block would lower the total number of initiation-competent ribosomes reaching uORF1under both normal or stress conditions, and thus affect the number of post-termination 40S ribosomes scanning downstream of uORF1’s stop codon. Further, reinitiation at uORF1 would be reduced under stress, as low TC levels hinder TC recruitment and production of a scanning competent PIC after termination from the start-stop. Initiation-incompetent ribosomes skipping the AUG of uORF1 would then be given more time to become initiation-competent en route to the AUG of uORF2, explaining its observed translation even under stress (**Figure 5A**) by a mechanism described elsewhere (23).

Translation of the wild-type reporter - i.e. the construct including the start-stop, uORF1, and the uORF2 fused to the firefly reporter - decreased under thapsigargin compared to normal conditions, but was still translated, albeit at lower levels (**Figure 5D**). Mutating the start-stop resulted in higher expression of the firefly reporter compared to the wild-type sequence, under both normal and stress conditions, again with reduced levels under stress (**Figure 5D**). The result suggests that the start-stop acts as a road-block which reduces ribosome flow along the 5’ UTR and lowers translation of uORF2 under both normal and stress conditions.

Examining these data more closely, we noticed a significant difference in the stress-induced drop in translation of uORF2, i.e. its bypass, as measured by the reduction of firefly signal under normal to stress conditions: stress-induced bypass of the second uORF was ∼4 times larger in the wild-type construct compared to the start-stop mutant (∼12% vs. ∼3%, p-value < 0.001, right panel, **Figure 5D**), after accounting for stress induced overall changes in translation as estimated by the renilla luciferase control. The renilla luciferase control decreased with stress due to the global decline in initiation during the stress response, but was not affected by the presence or absence of the start-stop (**Figure S5G**).

These results suggested a model in which, under normal conditions and in the absence of the start-stop, like in the mouse *Atf4* 5’ UTR, more ribosomes reach uORF1 than in the presence of the start-stop (**Figure 5E**). Both in the presence and in the absence of the start-stop, most ribosomes will reinitiate at uORF2 upon termination from uORF1 due to high TC abundance. However, as the start-stop lowers ribosome flow, it would also lower translation of uORF2, i.e. render the system more economical, in particular under normal conditions. Under stress, the start-stop promotes by-pass of uORF2 while still allowing its translation as observed by us and others (26, 66, 67), therefore supporting ATF4 induction. The precise interplay of the start-stop with the uORFs and other cis-acting elements is discussed elsewhere (23).

Finally, we tested whether our interpretation of the experimental observations could be matched by a simple mathematical model that reflected different regulatory scenarios (**Figure S5H, S5I**). To do so, we incorporated the quantitative measurements into the model and conducted deterministic simulations to examine the impact of the start-stop on ATF4 translation. In the model, we based a sigmoidal dependence of reinitiation at uORF2 on the levels of phosphorylated eIF2alpha, according to a Hill-Langmuir relationship of cooperative binding. We then examined the increases in reinitiation at the CDS as a result from lowering translation of uORF2. The simulation showed that even only 10% less overall reinitiation at uORF2 due to a start-stop reducing ribosome flow - but at the same overall availability of the ternary complex - would increase ATF4 translation under stress several fold compared to a model without the start-stop (**Figure S5H, S5I**). The model also confirmed that within biologically relevant ranges of phospho-eIF2alpha and *ATF4* mRNA levels, the start-stop is capable of mitigating the energy-demanding translation of uORF2, while ensuring even higher ATF4’s induction rate under stress.

## Discussion and conclusion

We present an in-depth characterization of start-stop elements in mammalian cells, a unique group of translation regulatory sequences that occur in the 5’ UTR of >1, 400 human genes and are evolutionarily conserved. Start-stop elements do not involve elongation, and are therefore distinct from open reading frames. We demonstrate that the impact of start-stops go beyond what has been described for individual genes (18, 21) or for a specialized case in plants (13, 17).

Start-stops often function in conjunction with other elements, e.g. uORFs, and share some of their characteristics, e.g. the repression of downstream translation. However, we demonstrate that start-stops are more than just ‘extreme’ uORFs and have distinct features, e.g. their ribosome retention that goes beyond what is expected based on initiation rates. By definition, start-stop elements have suboptimal nucleotides at the +4 and +5 initiation sites, i.e. a scanning 48S PIC is much less likely to initiate at a start-stop than at uORFs with an optimized initiation site (e.g. with a GCG as the second codon). We show that despite this difference, start-stops appear to retain the ribosome - once it initiated - much more than a uORF. This ribosome stalling at a start-stop drastically reduces the number of 40S subunits scanning downstream of the element (“ribosome flow”) and reduces downstream translation more so than in highly initiating uORFs.

Ribosome stalling on a start-stop does not appear to trigger NMD and even seems to slightly stabilize the transcript. This feature is another difference to uORFs where a delayed ribosome can drastically lower transcript half-life (64). A possible explanation for this difference is that translation of long uORFs can cause ribosome collisions which is a known trigger for NMD (71). Ribosomes on a start-stop cannot cause ribosome collisions, because of the absence of translocation which in turn prevents formation of two fully assembled ribosomes.

We hypothesized that ribosomes stall on start-stops either through incomplete termination which renders the 80S ribosome to be ‘stuck’ in a simultaneous initiation-termination state, or termination followed by partial recycling and rejoining of the 60S subunit (**Figure 4A**). Several lines of evidence suggest that the latter model is more likely. First, *in vitro* single-molecule analysis of start-stop elements suggested that they recruit release factors with the same kinetics as stop codons of longer open reading frames (72), not supporting possible defects in termination. Our analysis of A-site occupancy confirmed these findings: reads positioned to start-stops suggested an occurrence of occupied/empty A-sites at a frequency similar to an ‘average’ of the A-site occupancy found on the start or stop codon of the CDS.

Second, our analysis of the 40S data showed that start-stop elements attracted a comparatively large number of 40S reads, larger than that found on the start or stop codon of a CDS or uORF. The result supports the partial recycling model where the 40S ribosome with the Met-tRNA_i_^Met^ is still bound at the P-site and base-paired with the start codon even after termination. It is also consistent with findings that the 40S subunit lingers upon termination due to a delay between the first and second ribosome recycling steps (12, 73). Termination and partial recycling followed by re-joining of the 60S subunit provides a direct mechanism for the 80S ribosome being kept on the start-stop.

Start-stop elements have been discussed in the context of ribosome recycling and reinitiation (74), specifically with respect to the DENR/MCTS1 proteins (75, 76) which form a complex that triggers the release of deacylated tRNA from post-termination 40S ribosomes. Some work suggests that DENR/MCTS1 are needed for continuous scanning and re-initiation downstream of start-stop containing 5’UTRs (21), while recent work on yeast orthologs of the DENR/MCTS1 complex does not support this view (24). While indeed some of our start-stop containing genes overlap with DENR/MCTS1 targets identified in other work (**Table S10**), the DENR/MCTS1 function is linked to re-initiation *downstream* of the start-stop element in mammalian genes. Therefore, it does not explain the prevalence of footprints on start-stops and the apparent ribosome halt that we observe.

As start-stop containing genes frequently occur in transcription factors and signaling molecules which are often dosage-sensitive, one function of the start-stop might be to prevent harmful overproduction of these genes which could interfere with their proper functioning. Further, suppression at the level of translation also allows for precise subcellular localization of the mRNA prior to translation, as well as an increased speed of expression induction. In addition, as start-stop elements are very efficient in suppressing downstream translation, start-stops (rather than uORFs) might allow for the non-translated transcript to be stored, e.g. in stress granules. Supporting this idea, we observed a significant overlap between start-stop containing genes and transcripts enriched in stress granules (p-value < 0.01, **Table S10**)(77), e.g. *NFIA* and *DROSHA* shown in **Figure 1B**.

To illustrate the regulatory potential of start-stop elements, we identified examples where the inclusion and exclusion of start-stop elements in transcripts could modulate translation across tissues and conditions, e.g. USP1 and ZNF711 (**Figure S3C, S3D**). In addition, we hypothesized that mutations disrupting start-stops might offer a new mechanistic explanation for translation-based pathological phenotypes. A loss-of-function mutation in the start-stop would de-repress translation of a gene, leading to aberrant expression levels. Indeed, we identified example genes with naturally occurring sequence variants, e.g. single nucleotide polymorphisms (SNPs), located within or next to the start-stop. For example, we identified a SNP disrupting the start-stop of *PTPN11* (https://www.ncbi.nlm.nih.gov/snp/rs1280682918). Its 5’ UTR contains a start-stop but no other predictable translation regulatory sequences. *PTPN11* encodes the SHP-2 protein which participates in RAS/MAPK and PI3K signaling.

PTPN11/SHP-2 overexpression enhances, amongst others, progression of liver cancer (78). The start-stop disrupting SNP primarily occurs in the East Asian subpopulation where liver cancer is frequent (79).

Another example is represented by *UHMK1*, a serine/threonine RNA-binding protein kinase that promotes cell cycle progression and has been implicated in perturbations of mRNA export and translation leading to metabolic reprogramming and drug resistance in melanoma (80). We identified a SNP changing the guanidine immediately upstream of the start-stop in *UHMK1*’s 5’ UTR and therefore disrupting its Kozak sequence (https://www.ncbi.nlm.nih.gov/snp/rs553959593). Both the start-stop and an upstream uORF show higher ribosome occupancy compared to the AUG of the *UHMK1*’s coding region, suggesting translation repression of the gene. As *UHMK1* overexpression promotes colorectal cancer cell growth and drug resistance (81), derepressed *UHMK1* translation might contribute to the disease. Consistent with this hypothesis, the SNP is prevalent amongst African Americans and colorectal cancer is more common in this subpopulation (82).

Finally, we illustrated how a start-stop element expands the repertoire of translation regulation in the cell at the example of *ATF4*. In this gene, the start-stop does not function to reduce base-line ATF4 expression levels, as this task is performed by the uORF1-uORF2 system. We demonstrated that ribosome stalling at the start-stop element can reduce wasteful production of the uORF2 peptide and, at the same time, support ATF4 inducibility under stress.

In sum, start-stops represent a hitherto underappreciated element in the 5’ UTR that seems to control translation of many key regulators in humans. The lack of the elongation step in start-stops creates a non-canonical mechanism which delays or even stalls ribosome progression much more so than in the conceptually related uORFs. Therefore, start-stop elements drastically reduce the ribosome flow along the 5’ UTR without production of a peptide. If positioned near the next downstream initiation codon, start-stops reduce - similar to uORFs - the probability of reinitiation. In contrast to ribosome stalling within a coding region, which often leads to transcript turnover (83), start-stop elements slightly stabilize the transcript. This property might support localization and/or storage of the transcript until its translation is desired. Since start-stops often occur in combination with uORFs and other elements in the same 5’ UTR, they seem to represent an important part of complex regulatory systems that fine-tune translational control under various environmental conditions.

## Acknowledgments

We would like to thank Dr. Neville Sanjana for providing HAP1 cells and lentiCRISPRv2 plasmid and helpful discussions regarding CRISPR experimental design. This work was supported by the US National Institutes of Health 5R35GM127089 and Chan Zuckerberg Initiative (to CV); the Czech Science Foundation Grant of Excellence in Basic Research 19-25821X and CZ.02.01.01/00/22_008/0004575 RNA for therapy by ERDF and MEYS and the Praemium Academiae grant provided by the Czech Academy of Sciences (L.S.V.); the Fonds de Recherche du Québec – Santé Senior Award (283444) (to IT); and The Swedish Research Council and the Wallenberg Academy Fellow program (to OL).

## Author contributions

Conceptualization: JR, SH, MH, IT, LSV, CV

Methodology: JR, SH, SW, MPM, JL, XG, HZ, SM, VH, AH, LSV, CV

Analysis: all authors

Visualization: JR, SH, SW, MPM, MP, ZG, AL, KS, CV

Funding: LSV, CV Writing: JR, LSV, CV

Review & editing: all authors

