## Supplementary Notes for "Regulatory start-stop elements in 5’ untranslated regions pervasively modulate translation"

### Figure S1.

**A.** Graphs show ribosome footprint counts at each position is calculated across all start-stop (n=1,417), 9 nucleotide uORFs (n=1,331), and control genes (n=1,837), and the start and stop codon of the main ORFs (n=19,514), respectively. Average ribosome footprint was calculated as the total count at each position divided by the number of genes containing the respective elements. Data were taken from (1).

**B.** Ribosome occupancy was counted at the -12 position with respect  to the AUG of the occupied 9-nucleotide uORFs, and  plotted based on  the type of middle codon. The ribosome occupancy is the average read count, taken from data in (1).

nt - nucleotides

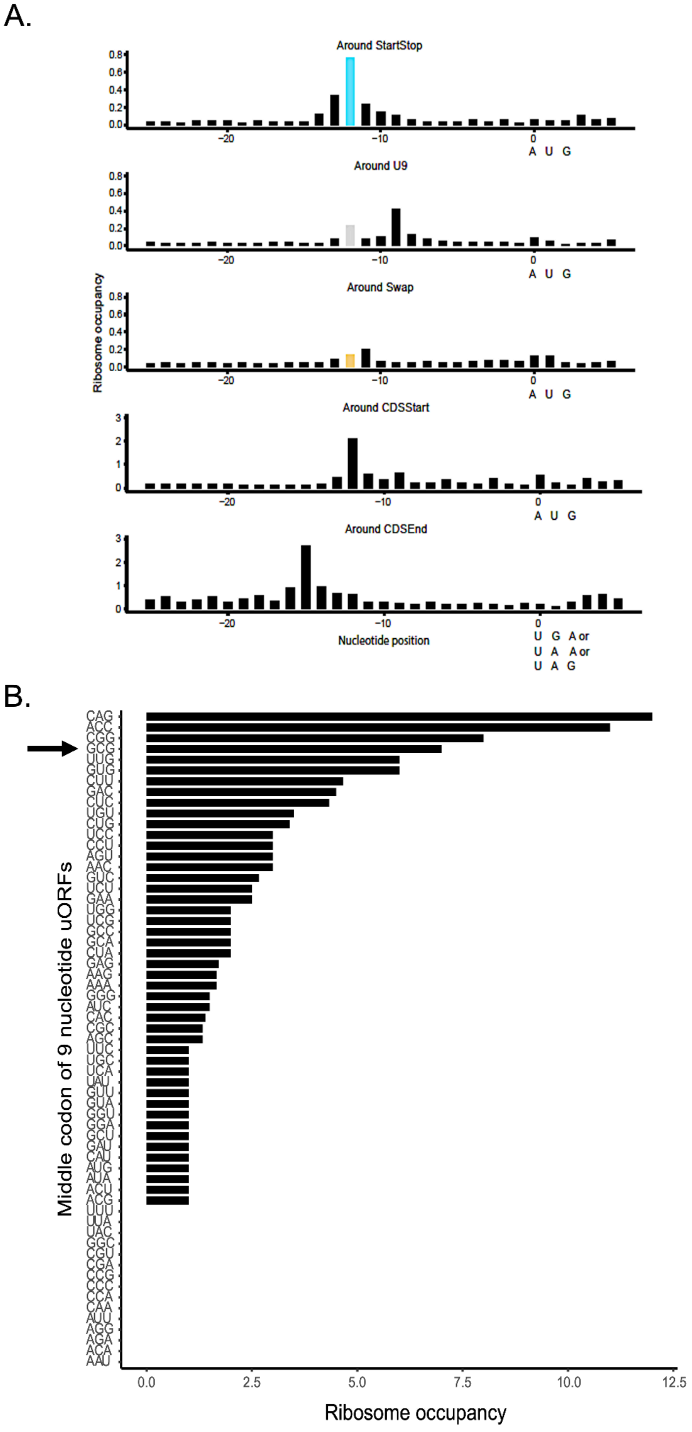

### Figure S2.

**A.** CDS translation efficiency of genes (n=312) with occupied start-stops compared to CDS translation efficiency of genes with occupied control sequences (n=121). Data on CDS translation efficiency was taken from (2). Graphs show average and standard error of the mean. We performed t-tests to determine the significance of the differences. * - P < 0.1; ** - P < 0.01; *** - P < 0.001. RPF - ribosome protected fragment

**B.** We used the GWIPS-vis browser (https://gwips.ucc.ie/) to assess ribosome occupancy at the start-stop in the D. melanogaster Mothers against dpp (Mad) gene. Coverage of this region was downloaded and plotted from S2 cell lines (3). The transcript is located on the reverse strand of chromosome 2L; genomic coordinates (BDGP R5/dm3) are denoted on the x-axis. The start-stop location is indicated by the dashed line.

**C.** Nucleotide frequencies for all four groups from the core set, complementing **Figure 2** in main text.

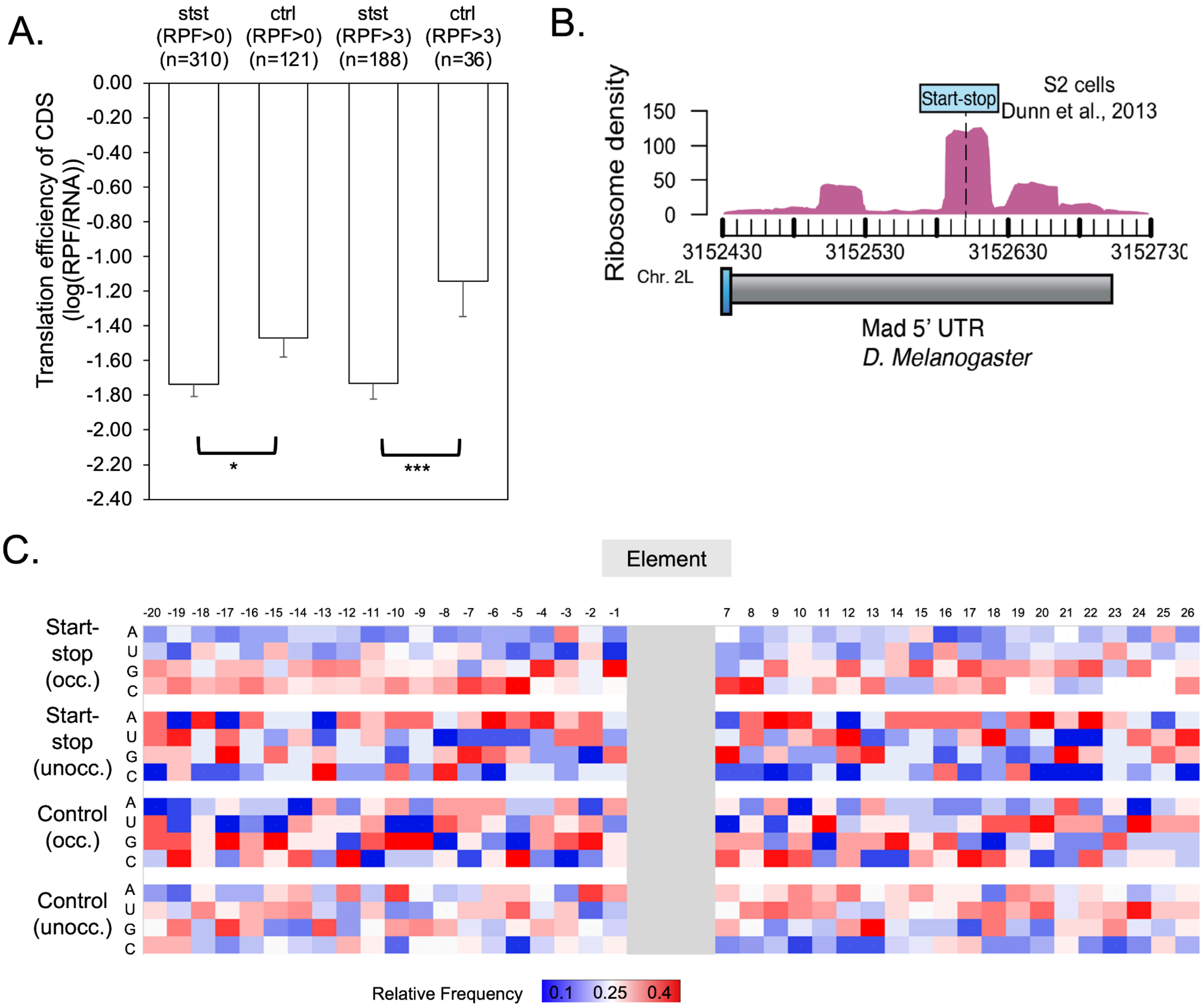

### Figure S3.

**A./B.** Examples of alternative transcript variants that include/exclude a start-stop through alternative exon usage: **A.** USP1 and **B.** ZNF711. For USP1, the start-stop is located on the most 5' exon which is specifically used for variant 1. ZNF711 contains three start-stops (stst1 to stst3) which occur in five transcript variants with different exon structures. For simplicity, other elements, e.g. uORFs are omitted from the diagrams. The start-stop is indicated by red arrows and the exon containing the start-stop is marked by red brackets. Shown are screenshots from gnomAD (https://gnomad.broadinstitute.org/) showing transcript variants and their tissue-specific relative expression. The data and original plots for USP1 and ZNF711 can be found here (https://gnomad.broadinstitute.org/gene/ENSG00000162607?dataset=gnomad_r4) and here (https://gnomad.broadinstitute.org/gene/ENSG00000147180?dataset=gnomad_r4).

**C.** RNA levels measured by RT-qPCR of start-stop genes with 5' UTR containing wild-type start-stop sequence (WT) and AUGUAG->UAGUAG or AUGUAG->AUUGAG, respectively (St-st Mut) measured in either HEK293T or HeLa or both cells. Plots below show the luciferase data for the same constructs. All results are relative to wild-type (set to 1).

Constructs are: MORF4L1 WT and AUGUAG->UUGUAG (St-st Mut), SLC39A1 WT and AUGUAG->UUGUAG (St-st Mut) and PSPC1 WT and AUGUAG->AUUGAG (St-st Mut). All experiments had been performed in triplicate. Graphs show the average and standard error of the mean. We performed t-tests to determine the significance of the differences. * - $P < 0.1$; ** - $P < 0.01$; *** - $P < 0.001$.

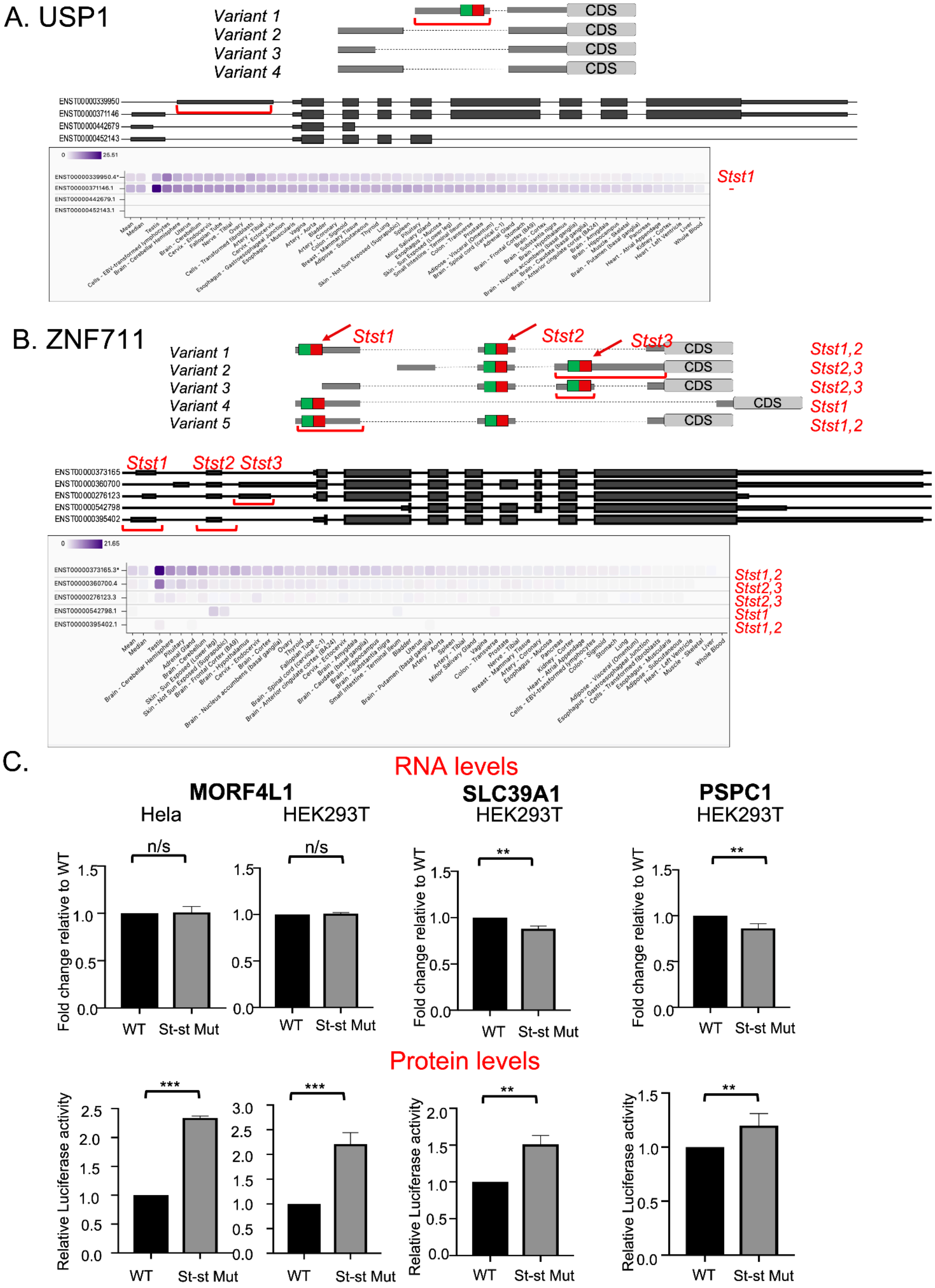

A. USP1
Variant 1
Variant 2
Variant 3
Variant 4
CDS
ENST00000339950
ENST00000371146
ENST00000442679
ENST00000452143
Stst1
B. ZNF711
Variant 1
Variant 2
Variant 3
Variant 4
Variant 5
Stst1
Stst2
Stst3
CDS
Stst1,2
Stst2,3
Stst2,3
Stst1
Stst1,2
ENST00000373165
ENST00000360700
ENST00000276123
ENST00000542798
ENST00000395402
Stst1
Stst2 Stst3
C.
RNA levels
MORF4L1
Hela
HEK293T
SLC39A1
HEK293T
PSPC1
HEK293T
Fold change relative to WT
n/s
**
WT
St-st Mut
Protein levels
Relative Luciferase activity
***
**

### Figure S4.

**A.** Expansion of start-stop to a longer uORF containing 5 codons (u21, GCG codon) in a gain-of-function construct similar to that described for **Figure 4.** The diagrams explain the constructs used. We inserted either start-stop ('Start-stop') or 21 nucleotide uORF ('uORF', defined as AUG(GCG)5UAG) sequences into the ACTB 5' UTR used for the luciferase reporter. The unmodified ACTB 5' UTR represents the wild-type which all measurements had been normalized to. It has a length of 84 nucleotides. The start-stop or uORF were positioned either 77 or 30 nucleotides upstream of the luciferase CDS to mark 'near cap' or 'near CDS' positions, respectively. The bar plot shows the protein level measured by luciferase assays. All results are relative to the wild-type ACTB 5'UTR set to 1. Experiments were performed in triplicate. The plot shows means and standard error of the mean.

**B.** The graph is similar to that shown in **Figure 4** with data taken from (4) which screened a library of randomly designed nucleotide sequences for their effect on translation and transcript stability, respectively. We downloaded the data and extracted start-stop, 9 nucleotide uORF, and control sequences from the data with the following criteria: start-stop sequences are "AUGUGA" or "AUGUAA" or "AUGUAG", 9 nucleotide uORF sequences start with "AUG" and end with "UGA" or "UAA" or "UAG", with one of the 61 sense codons in the middle, and control sequences are "AUUGGA" or "AUUGAA" or "AUUGAG". We required no occurrence of "AUG" up- or downstream of a start-stop or control sequence, to avoid the confounding effect of potential uORFs. We further obtained constructs containing 9 nucleotide uORFs with a good Kozak context: those with the middle codon starting with "GA" or "GC". Graphs show means and standard error of the mean.

Note that the graph shows translation *inefficiency*, i.e. the ratio of mono- vs. polysome occurrence of the respective transcripts. We performed t-tests to determine the significance of the differences. * - $P < 0.1$; ** - $P < 0.01$; *** - $P < 0.001$.

**C.** Graphs show ribosome footprint counts at each position is calculated across all start-stop (n=1,417), 9 nucleotide uORFs (n=1,331), and control genes (n=1,837), and the start and stop codon of the main ORFs (n=19,514), respectively. Average ribosome footprint was calculated as the total count at each position divided by the number of genes containing the respective elements. Reads were split into short (20-22 nt) and long (27-30 nt) reads, corresponding to empty and occupied A-site-containing ribosomes, respectively. Data were taken from (1). nt - nucleotides

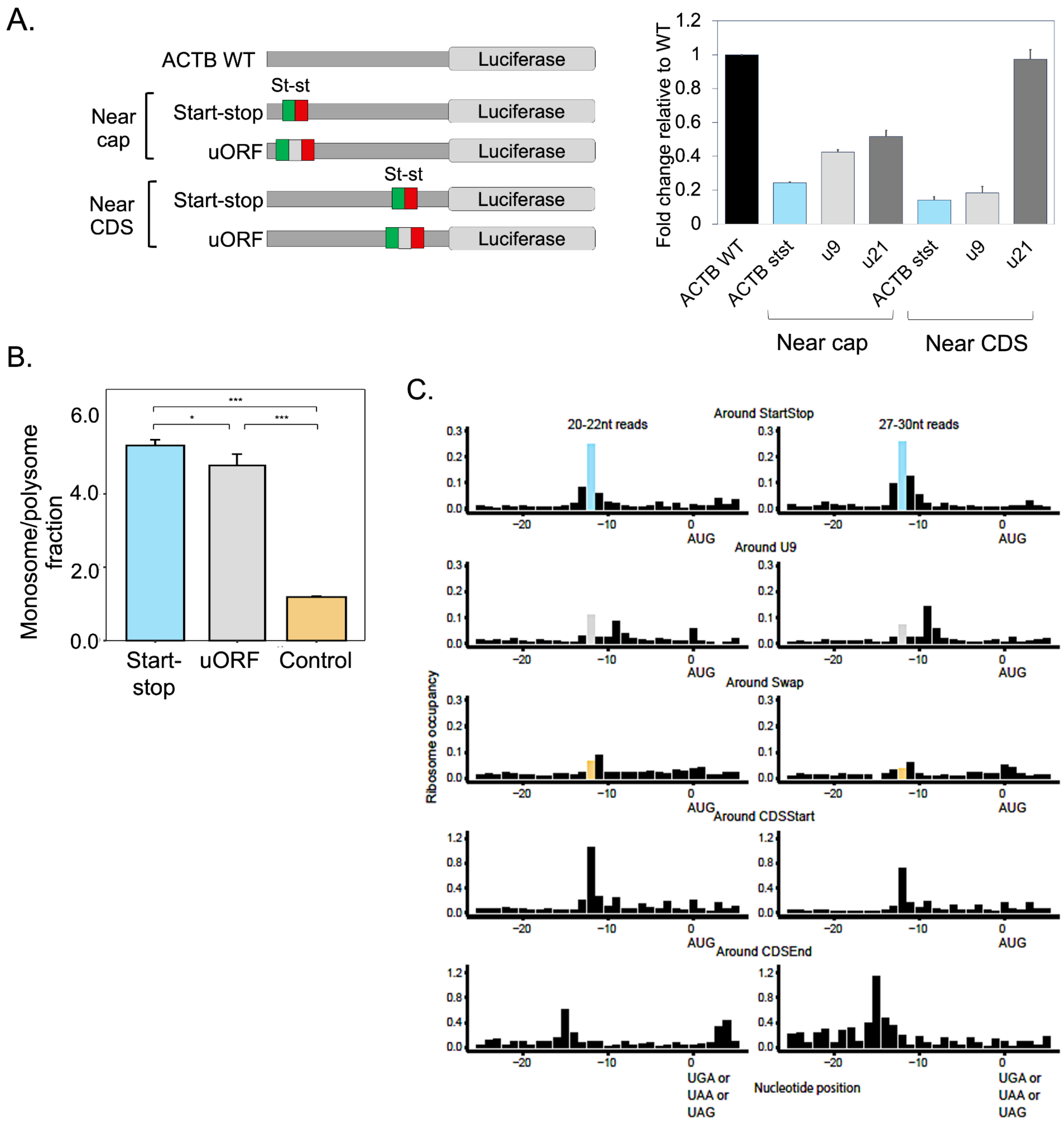
A.
ACTB WT
Luciferase
Near cap
St-st
Start-stop
uORF
Near CDS
Fold change relative to WT
ACTB WT
ACTB stst
u9
u21
B.
Monosome/polysome fraction
Start-stop
uORF
Control
C.
Around StartStop
20-22nt reads
27-30nt reads
Around U9
Around Swap
Around CDSStart
Around CDSEnd
Ribosome occupancy
AUG
UGA or UAA or UAG
Nucleotide position

### Figure S5.

**A.** Genomic alignments of ATF4's 5' UTR start-stop region from three mammalian species using the UCSC Genome Browser (https://genome.ucsc.edu/index.html)(5–11). Below each nucleotide we display the phyloP-based vertebrate conservation scores. Scores were measured using the phyloP method from the PHAST package. Evolutionary conservation scores from 100 vertebrate species were compared within a 30 nt region centered on the start-stop.

**B.** Ribosome density (total ribosome protected fragments) around the start-stop in the 5' UTR of ATF4 is plotted for different human cell lines, using data from four published datasets. Top to bottom: HeLa (2), HEK293 (11) , HAP1 (12), and iPSC cell lines (13). The citation for each and the method used during sample collection is noted on the right side of each profile (cycloheximide, No drug, harringtonine). The position (GRCh38/hg38) in Chr. 22 is denoted along the x-axis, and the location of the start-stop is indicated by the dashed lines.

**C.** Representative Western blots of ATF4 and loading control for CRISPR-edited HAP1 cells before and after treatment with thapsigargin (Tg, 1uM). Below are ATF4 protein levels (left n = 3, right n = 2) normalized to the loading control (% of max).

**D.** Luciferase activity for start-stop only and start-stop overlapping with main ORF. Experiments were conducted as described before and in triplicate.

**E.** 18S rRNA levels for Start-stopWT and delta-Start-stop HAP1 cell lines before and after treatment with ER stressor thapsigargin (Tg, 1uM); measured by qPCR (n = 3 for each time point).

**F.** ATF4 mRNA levels after treatment with RNA polymerase II inhibitor (DRB, 5,6-dichloro-1-β-ribofuranosyl-benzimidazole). Right panel shows calculated half-lives in hours (h).

**G.** Activity of the internal control Renilla luciferase that is co-expressed from the same plasmid as reporters for uORF2 reinitiation assay.

Graphs **C-G.** show mean and standard error of the mean. We performed t-tests to determine the significance of the differences. * - $P < 0.1$; ** - $P < 0.01$; *** - $P < 0.001$.

**H**. The graphs show the output of the quantitative modeling to illustrate the start-stop's role in regulatory efficiency and sensitivity. To quantify the impact of the start-stop on the reinitiation of uORF2 during stress independently of global changes in initiation associated with eIF2 phosphorylation, we calculated the change in uORF2 bypass as:

$$\Delta\text{uORF2 bypass}=\Delta\text{uORF2 initiation}-\Delta\text{global initiation}$$

Next, we incorporated parameters measured in our experiments to verify the proposed model. We compared a system similar to ATF4 from mouse that only relied on bypass of the second uORF for translational induction ('uORF-only') to a system in which a start-stop preceded the uORFs as found for human ATF4 ('Start-stop + uORF') to a system that only relied on uORFs but doubled transcription ('2x transcription + uORF'). We simulated values for ATF4 mRNA, uORF2 peptide, and ATF4 protein levels for each system under three differing conditions (from left to right): 1) cells reaching steady state under control (normal) conditions, 2) cells transitioning from control conditions to stress, and 3) cells experiencing acute stress followed by recovery.

Under control conditions, none of the models produced ATF4 protein, as is observed in cultured cells. The model in which transcription was doubled reached the highest mRNA levels at steady state, but also produced twice as much peptide from the second uORF. The model including both the start-stop and uORFs exhibited an increase in ATF4 mRNA levels relative to the uORF-only model, due to decreased mRNA decay. Importantly, this start-stop mediated mRNA stabilization did not increase wasteful uORF2 translation as the start-stop also reduced the flow of scanning ribosomes within the 5' UTR. However, the most striking difference among the models was the greater stress-inducibility of ATF4 protein of the model including uORFs and a start-stop compared to other models: a small decrease (~10%) in the probability of initiating at uORF2 due to an upstream start-stop resulted in >400% increase in induction of the CDS. This finding aligned with our observations in ribosome profiles of cells undergoing ER stress, where small decreases in ribosome occupancy at uORF2 coincided with large increases in CDS occupancy. Additionally, under acute stress, the uORF-only and the uORFs+start-stop models produced ATF4 protein levels that were similar to those observed for delta-Start-stop and Start-stop wild-type cell lines, respectively, indicating that indeed, the uORFs are essential and sufficient for ATF4 translation under stress, while the start-stop increases the sensitivity of induction.

**I**. Graphs display hypothetical values based on the assumption that all scanning ribosomes along the ATF4 transcript will reinitiate at either the second uORF (uORF2) or the CDS and that 99% of ribosomes reinitiate at uORF2 under normal conditions. Lines and bars correspond to uORF2 (in grey) and CDS (in blue). Panel 1 demonstrates sigmoidal dependency between uORF2 reinitiation and the levels of phosphorylated eIF2a (P-eIF2a) according to the Hill-Langmuir relationship of cooperative binding, and the corresponding increases in CDS reinitiation. The dashed line indicates the level of translation observed in HeLa cells upon 8 hours of tunicamycin treatment. The levels of translation were based on ribosome occupancy on uORF2 which were set to 100% at time 0. Panel 2 focuses on the portion of the graph corresponding to experimental conditions used here (zoom-in). Panel 3 shows the log2(fold change) of uORF2 and CDS ribosome occupancy that would result from a 10% reduction in total ribosome occupancy (relative initiation levels) at either uORF2 or the CDS based on reduced ribosome flow due to the start-stop. In Panel 4 the arrows indicate two possible ways in which the ATF4 system would be modified through the inclusion of the start-stop, compared to the model without a start-stop in mouse: 1) lowering the basal uORF2 translation to reduce waste. 2) increasing the sensitivity of CDS induction.

Tg, thapsigargin; st-st, start-stop. *P < 0.05, **P < 0.01; paired t-tests.Graphs show mean and standard error of the mean.. St-St, start-stop.

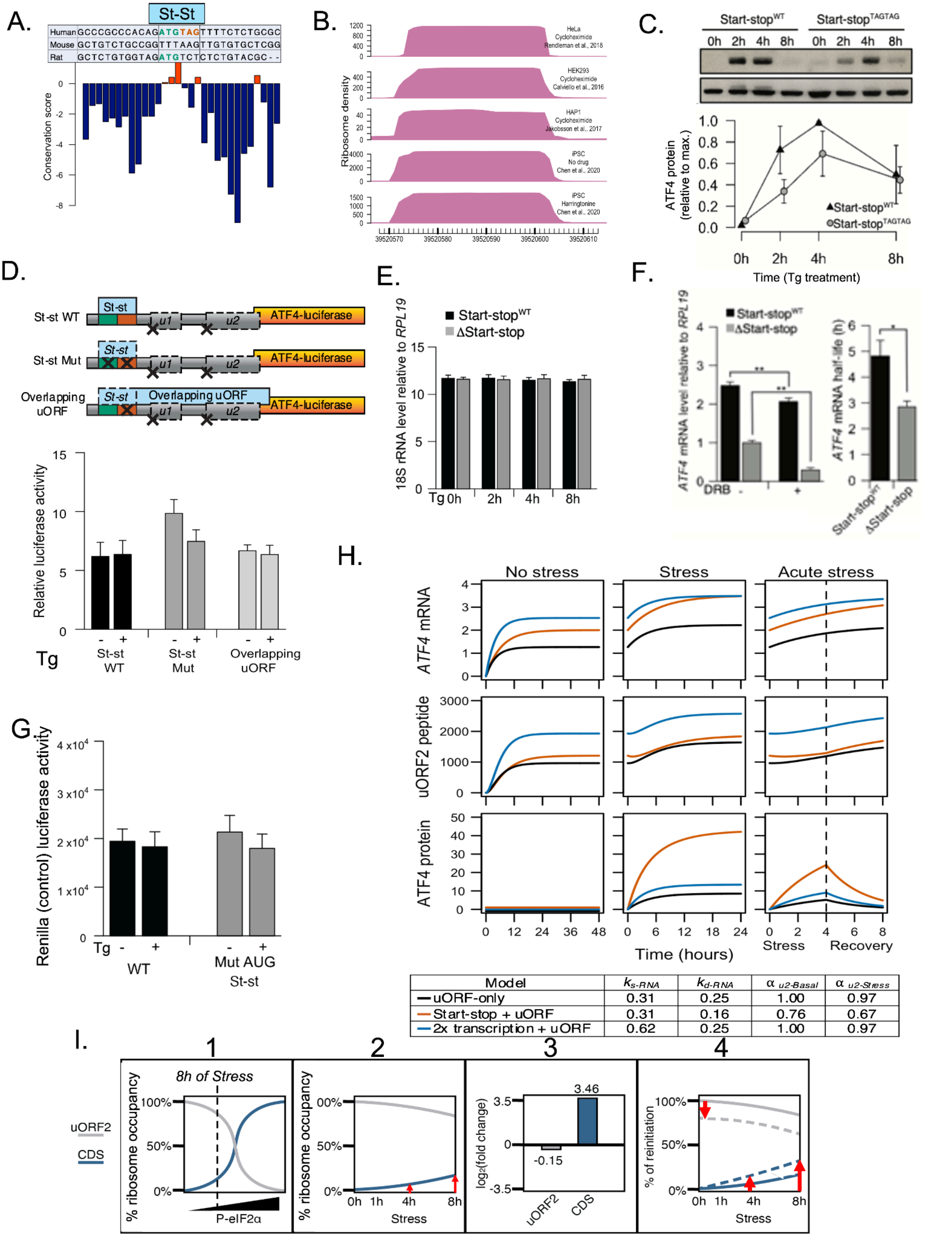


| Model | $k_{s\text{-}RNA}$ | $k_{d\text{-}RNA}$ | $\alpha_{u2\text{-}Basal}$ | $\alpha_{u2\text{-}Stress}$ |
|---|---|---|---|---|
| uORF-only | 0.31 | 0.25 | 1.00 | 0.97 |
| Start-stop + uORF | 0.31 | 0.16 | 0.76 | 0.67 |
| 2x transcription + uORF | 0.62 | 0.25 | 1.00 | 0.97 |
